# Targeting ALOX5/LTA4H driven granuloma caseation as a host-directed strategy for control of TB associated lung damage

**DOI:** 10.1101/2025.02.27.640170

**Authors:** Thabo Mpotje, Kerishka Rajkumar-Bhugeloo, Denelle Moodley, Kievershen Nargan, Tamia KJ Lawrence, Kimone L Fisher, Setjie W Maepa, Kershnee Thambu, Kapongo Lumamba, Pumla Majozi, Antony M Rapulana, Suraj Parihar, Threnesan Naidoo, Rajhmun Madansein, Hlumani Ndlovu, Alasdair Leslie, Siamon Gordon, Adrie Steyn, Mohlopheni J Marakalala

**Affiliations:** Africa Health Research Institute, Durban, South Africa; School of Laboratory Medicine and Medical Sciences, Nelson Mandela School of Medicine, University of KwaZulu-Natal, Durban, South Africa; Institute of Infectious Disease and Molecular Medicine, Division of Chemical & Systems Biology, University of Cape Town, Cape Town, South Africa; Institute of Infectious Disease & Molecular Medicine, Department of Pathology, University of Cape Town, Cape Town, South Africa; Department of Forensic and Legal Medicine, Walter Sisulu University, Mthatha, South Africa; Department of Cardiothoracic Surgery, University of KwaZulu Natal, Inkosi Albert Luthuli Central and King DinuZulu Hospitals, South Africa; Chang Gung University, Taoyuan, Taiwan; Department of Cellular pathology, Sir William Dunn School of Pathology, University of Oxford; Division of Infection and Immunity, University College London, London, United Kingdom; Department of Microbiology, University of Alabama, Birmingham, AL 35294, USA; Centers for AIDS Research and Free Radical Biology, University of Alabama at Birmingham, Birmingham, AL 35294, USA; South African Medical Research Council Centre for Tuberculosis Research, Division of Molecular Biology and Human Genetics, Faculty of Medicine and Health Sciences, Stellenbosch University, Cape Town

**Keywords:** Inflammation, ALOX5, LTB4, LTA4H, caseous TB granuloma, tuberculosis, macrophages, histopathology, HDTs

## Abstract

Tuberculosis remains one of the major global health challenges with those previously exposed presenting persisting pulmonary dysfunction. Therefore, improved control measures are still required to prevent or reduce the TB-induced immunopathology in affected individuals. Host directed therapies that target host factors, have been suggested to improve on current treatments against TB. We previously reported a potential role for proteins which metabolize the arachidonic acids during TB immunopathogenesis. The study, therefore, sought to demonstrate the potential role of these proteins in TB-induced immunopathology. Using immune-histopathological assays, the association of macrophage driven ALOX5 signaling with severe pulmonary immunopathology during TB was demonstrated. Furthermore, using both *in vitro* and *in vivo* assays, the data demonstrated the contribution of ALOX5 signaling to TB-induced granulomatous inflammation. Interception of the signaling pathway through clinically approved pharmaceutical inhibitors, resulted in reduced lung damage during the disease. The resolution of TB-induced lung pathology further contributed to increased bactericidal activity. Taken together, our data strongly suggest a role for macrophage driven inflammation via ALOX5 signaling which can be targeted for the development of therapeutic treatment to prevent exacerbated TB-induced pulmonary immunopathology.

## Introduction

Tuberculosis remains one of the major global health concerns contributing to over 10.6 million annual cases reported^1^. The disease is primarily driven by the development of a characteristic feature called granuloma formation which is mediated by the host immune cellular responses to the disease-causing pathogen^2,3^. Although the disease is curable with over 84% success rate, the affected individuals still suffer long lasting effects from the resultant pathology^1,4^. Furthermore, the pathology also contributes to reduced access to the site of infection by currently available TB drugs^5,6^. Because of this, TB therefore, contributes to a growing global burden of lung functions highlighting the need for earlier interventions to understand and prevent tissue damage, which contributes to diminishing lung function in affected individuals^4,7^.

Granulomas are a pathological hallmark of TB disease formed through cellular aggregates mediated initially by macrophage populations upon recognition and phagocytosis of the invading bacteria^8,9^. Other immune cell types, including dendritic cells and T cells, are recruited to the site of infection to boost the antimicrobial properties of macrophages against the bacteria^10,11^. Formation of the granuloma helps contain and neutralize the bacteria through antimicrobial properties driven largely by macrophages^8^. When the cells fail to control the infection, they undergo cell death. This leads to the development of a necrotic core of *Mtb*-infected alveolar macrophages surrounded by an inflammatory layer composed primarily of macrophages and an outer layer of a lymphocytic cuff^10–14^. Therefore, the caseous TB granulomas are characterized by macrophages and multinucleated giant cells forming the inner layer that is surrounded by T lymphocytes^15–17^. Failure of the granulomas to contain the bacteria contributes to the disease progression and clinical outcome^14,18^. Therefore, understanding the immune mechanism for the control of *Mtb* infection is vital to control tissue mediated pathology and thus clinical outcome.

Host directed therapies (HDTs) have recently been suggested as potential strategies to improve the current control for TB^19^. This is because, these are useful in exploiting host factors that modulate key inflammatory signals during disease progression^20^. The therapeutic approach offers a promising adjunct strategy to control TB and the resultant tissue damage, by modulating inflammatory responses^21^. For instance, a study by Mayer-Barber (2014) provided a proof of concept for host-directed therapies through manipulation of the eicosanoid network to improve resistance against *Mtb* infection^22^.

The eicosanoid network is triggered when immune cells, composed of polyunsaturated fatty acid, are activated by various stimuli leading to production of arachidonic acid (AA) through phospholipase A_2_^23^. Further metabolism of AA by metabolic pathways including arachidonate 5-lipoxygenase (5-LO) and the cyclooxygenase (COX) pathways lead to production of leukotrienes (LTs) and prostaglandins, respectively^24–26^. The LT family includes, LTB4, which is a known pro-inflammatory mediator that affects immune cells such as macrophages, polymorphonuclear leukocytes (PMN), natural killer (NK) cells and dendritic cells^27^. Studies have demonstrated the significance of LTs in mediating inflammatory responses during disease complications^27–29^. Recently, the ALOX5/LTA4H mediated pathway has been hypothesized to associate with TB-induced lung damage, suggesting a role for the signaling pathway during TB disease progression^30^. Our study, therefore, aimed to explore in detail the role of ALOX5/LTA4H signaling as a potential therapeutic strategy to manage TB induced lung damage.

## Results

### ALOX5 associates with increased immunopathology in human TB granulomas

We have previously utilized laser capture microdissection and MS-proteomics to identify protein signatures in human TB granulomas^30^. ALOX5, a protein that metabolizes arachidonic acid to produce lipoxins and leukotrienes, was predicted to be more abundant in the necrotic regions of caseous granulomas^30^. We sought to characterize the role of the ALOX5 pathway in TB pathogenesis (Figure S1a). Using immunohistochemistry, we stained for this protein on heterogeneous forms of human TB granulomas. ALOX5 was more abundant in the caseum compared to cellular granulomas (Figures S2a - f). Interestingly, the protein was uniquely enriched in the inflammatory borders inside the caseum (Figures 1A; S2g and h). To confirm the spatial association of ALOX5 with inflammatory lesions, we utilized immunofluorescence staining of caseous granulomas with varying degrees of damage/severity (Figures **1a** and **b**; Table 1). Consistently, there was an enrichment of ALOX5 in the inflammatory layer (region B) immediately bordering the caseum (Figures **1b** and **c**). This was confirmed using QuPath software (Figure S3), which revealed that there were more cells expressing the protein in the inflammatory layer (Figures **1c** and **d**). Low numbers of cells in the necrotic centre (region A) as well as the peripheral/outer cellular rings (region C) outside the caseum expressed the protein (Figures **1c** and **d**). Taken together the results confirm the association of ALOX5 with caseous lesions in TB granulomas.

**Figure 1:**
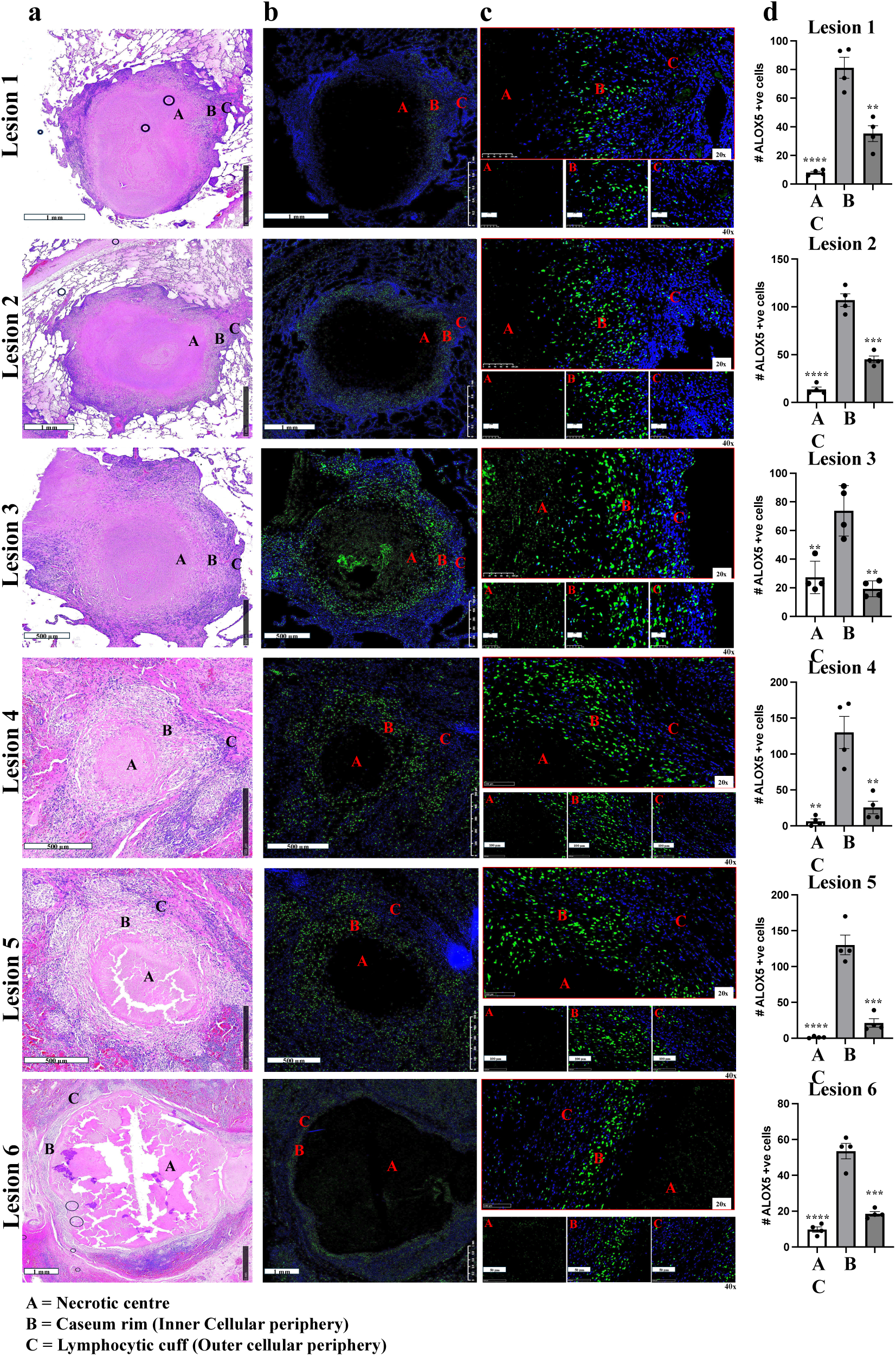
ALOX5 associates with increased immunopathology in human TB granulomas. **a**. Using lung tissue biopsies collected TB-induced lesions were screened and 6 granuloma types were identified, signifying the degree of damage using H&E staining. **b,** ALOX5 expression was quantified using immunofluorescence on each of the granuloma types. **c**, IF staining showing the spatial expression pattern of ALOX5 on the necrotic, inflammatory ring, and the lymphocytic cuff regions of a TB-induced granuloma. **d**, Quantification of ALOX5 expressing units (cells) using Qupath software (QuPath v0.5.0)^50^ in the necrotic, cellular, and the surrounding regions of a TB-induced granuloma. Error bars denote mean ± SEM. *P < 0.05; **P < 0.01; ***P < 0.001; Student’s t-test was used to measure the significance of the differences between groups.

**Table 1:** Characterization of TB-induced granuloma lesions.

| Granuloma | Description |
| --- | --- |
| Lesion 1: | Characterized by necrotic centre, and cellular periphery composed mostly of lymphocytes. |
| Lesion 2 | Characterized by central necrosis containing karyorrhectic debris and is surrounded by layer of activated macrophage as well as the lymphocytic cuff respectively. |
| Lesion 3 | Characterized by centre with karyorrhectic debris together with an infiltration of histiocytes staining positive for ALOX5. The centre is surrounded by a caseum rim and a lymphocytic cuff layer. |
| Lesion 4 | Characterized by necrotic centre that is surrounded by peripheral cellular ring containing epithelioid histiocytes (activated macrophages), and a lymphocytic cuff that is not well-formed. |
| Lesion 5 | Characterized by central necrosis with cavitating centre. The granuloma also has a cellular periphery containing activated macrophages epithelioid histiocytes, fibroblasts and a few multinucleated Langhans giant cells. Furthermore, the lesion also contains vessels in the fibrotic area, which hints at a healing fibrosis. |
| Lesion 6 | Characterized by necrotic centre surrounded by activated macrophage layer and lymphocytic cuff. |

### ALOX5 mediated signaling is largely expressed by inflammatory macrophage populations during TB

Given the unique spatial enrichment of ALOX5 in the inflammatory layer, we investigated the characteristic features of the granuloma regions including necrotic centre, inflammatory layer, and the lymphocytic cuff. Although the necrotic centre had few to no cellular populations (region A), we noted the inflammatory layer (region B) of the caseous granuloma mainly contained cell populations that resembled the myeloid cells, while the peripheral ring (region C) contained cells that resembled the lymphocyte populations (Figure **2a**). Taken together, we hypothesized that ALOX5 signaling was largely driven by the inflammatory macrophages during TB. To confirm this, immunofluorescence staining was used to demonstrate an increased abundance of CD68 positive cells (inflammatory macrophages) in the inflammatory layer of the caseous granuloma where the ALOX5 was also enriched (Figures **2b****.i** and **ii**; **2c** and **2d**). Further analysis confirmed unique cell populations where co-localization of CD68 and ALOX5 was observed (Figure **2b****.iii**). As predicted, we have confirmed that the ALOX5 signaling is largely expressed by the macrophage populations. Furthermore, cells are uniquely enriched in the inflammatory layer of caseous TB granuloma.

**Figure 2:**
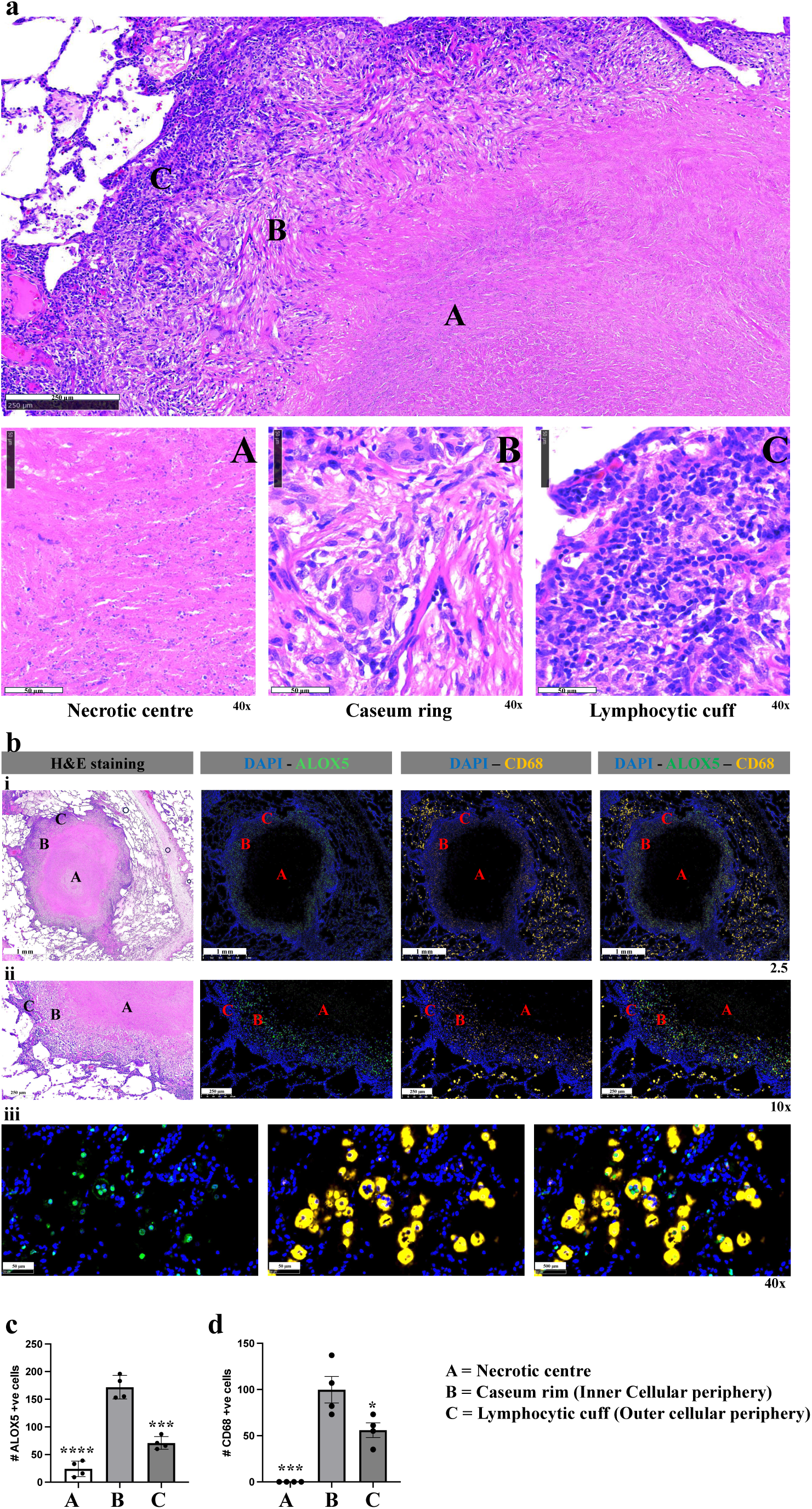
ALOX5 mediated signaling is driven largely by inflammatory macrophage populations during TB. **a**, H&E staining of a TB caseous granuloma to show the spatial recruitment of cells in the necrotic, inflammatory ring, and lymphocytic cuff regions of a TB-induced granuloma. **b**.**i**, Expression of ALOX5 together with CD68 were quantified using immunofluorescence on caseous TB granuloma to demonstrate their association. **b.ii**, Staining of ALOX5 and CD68 to demonstrate their spatial association in the same regions including the necrotic, inflammatory ring, and lymphocytic cuff. **b.iii**, **c, d**, Staining of ALOX5 and CD68 to demonstrate their co-localization within the same cell populations. Quantification of CD68 and ALOX5 expressing units (cells) using QuPath software in the necrotic, inflammatory ring, and lymphocytic cuff of a caseous TB granuloma. Error bars denote mean ± SEM. *P < 0.05; **P < 0.01; ***P < 0.001; Student’s t-test was used to measure the significance of the differences between groups.

### *ALOX5* is upregulated in blood of TB patients

The enrichment of ALOX5 within regions of high immunopathology in TB lungs suggested that the protein may be associated with the clinical disease. To determine the association of ALOX5 with clinical TB, we investigated the expression patterns of the gene in blood from healthy, latently infected individuals and TB patients. ALOX5 was upregulated in TB patients compared to both LTBi and healthy individuals (Figure **3a**). However, the expression levels of the gene were significantly higher in LTBi than healthy controls, suggesting ALOX5 may be associated with severity of the disease. Interestingly, ALOX5AP, a gene that activates ALOX5, was also upregulated in the blood of TB patients (Figure **3b**), indicating the importance of this pathway in TB pathogenesis. Using ELISA, we found that the plasma protein levels of ALOX5 were more abundant in TB than in LTBi patients (Figure **3c**). These results were corroborated by Western Blot analysis of the protein in PMBCs, where ALOX5 was detectable in higher levels in cells from TB patients compared to LTBi and healthy controls (Figure **3d**). The expression levels of COX 1 and 2, which mediate an alternative pathway of AA metabolism, in active TB participants were like those of healthy and LTBi individuals (Figure S1b). Therefore, the findings suggest that the ALOX5 pathway associates with TB in blood and that it may indicate clinical severity of the disease.

**Figure 3:**
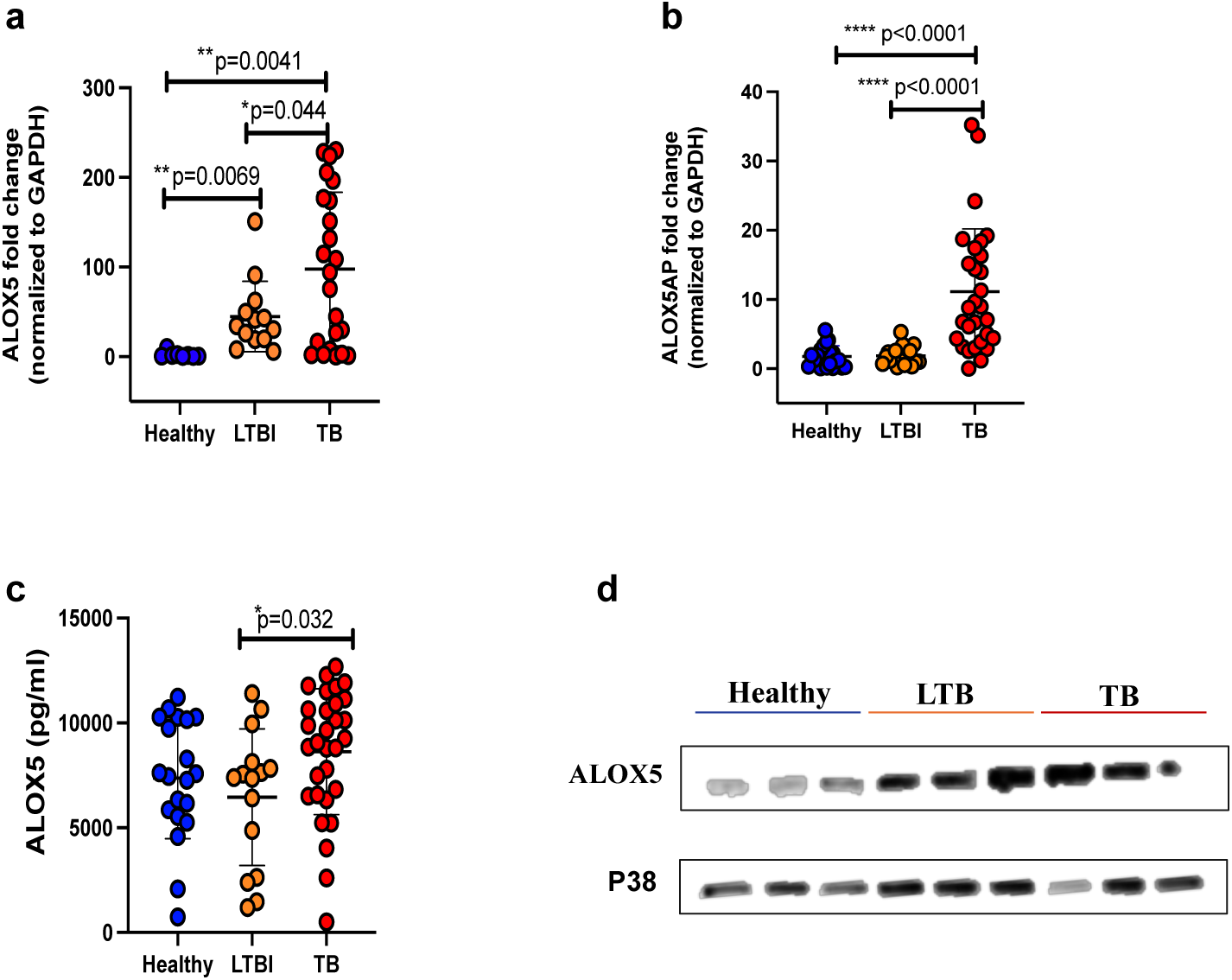
*ALOX5* is upregulated in blood of TB patients. **a,b,** Expression levels of ALOX5 and ALOX5AP genes were quantified using qPCR from RNA samples extracted from 20 healthy, 20 latent, and 30 active TB participants using primers specific for the genes and normalized to GAPDH. **c,d,** The expression levels of the ALOX5 were further confirmed at a protein level using ALOX5 specific ELISA kit, and western blot from healthy, latent, and active TB participants using anti-ALOX5 antibody in relation to p38 housekeeping protein. The sample sizes for qPCR and ELISA assays included 20 healthy, 20 latent, and 30 active TB participants. For western blot, the sample size was 3 healthy, 3 latent, and 3 active TB participants. Error bars denote mean ± SEM. *P < 0.05; **P < 0.01; ***P < 0.001; Student’s t-test was used to measure the significance of the differences between groups.

### ALOX5 activates an LTA4H/LTB4 mediated signaling axis in TB

ALOX5 drives arachidonic acid (AA) degradation leading to production of downstream lipid mediators, including lipoxins and leukotrienes, which may in turn contribute to inflammatory responses^31^. The association of ALOX5 with TB lung inflammatory lesions suggests that this protein may contribute to disease pathogenesis through induction of inflammatory leukotrienes. To explore this, we investigated inflammatory lipid mediators downstream of the ALOX5 signaling pathway. We measured leukotriene B4 (LTB4) in peripheral blood mononuclear cells infected with *Mtb*. The infection resulted in significant release of LTB4, which was reduced upon administration of the ALOX5 inhibitor, Zileuton (Figure **4a**). Furthermore, we observed a moderate increase in plasma LTB4: AA ratio in TB participants compared to healthy and LTBI groups (Figure **4b**). This suggests that ALOX5 mediates TB related inflammatory responses through degradation of AA to release its downstream lipid mediator, LTB4. We then sought to investigate the association of LTA4H, a hydrolase that catalyzes the final step in the LTB4 synthesis^32^, with the disease. Interestingly, LTA4H was significantly enriched in the whole blood samples of TB compared to LTBi and healthy participants (Figure **4c**), mirroring the phenotype observed with ALOX5. The plasma levels of LTA4H protein were also higher in TB participants compared to LTBi and healthy groups (Fig. **4d**), suggesting that this protein could be a marker of TB disease, as indicated by an AUC of = 0.82 and sensitivity and specificity of = 0.73 and 0.75, respectively (Figure **4e**). Together, these results indicate that ALOX5 association with TB may stem from its induction of inflammation through the LTA4H/LTB4 signaling axis.

**Figure 4:**
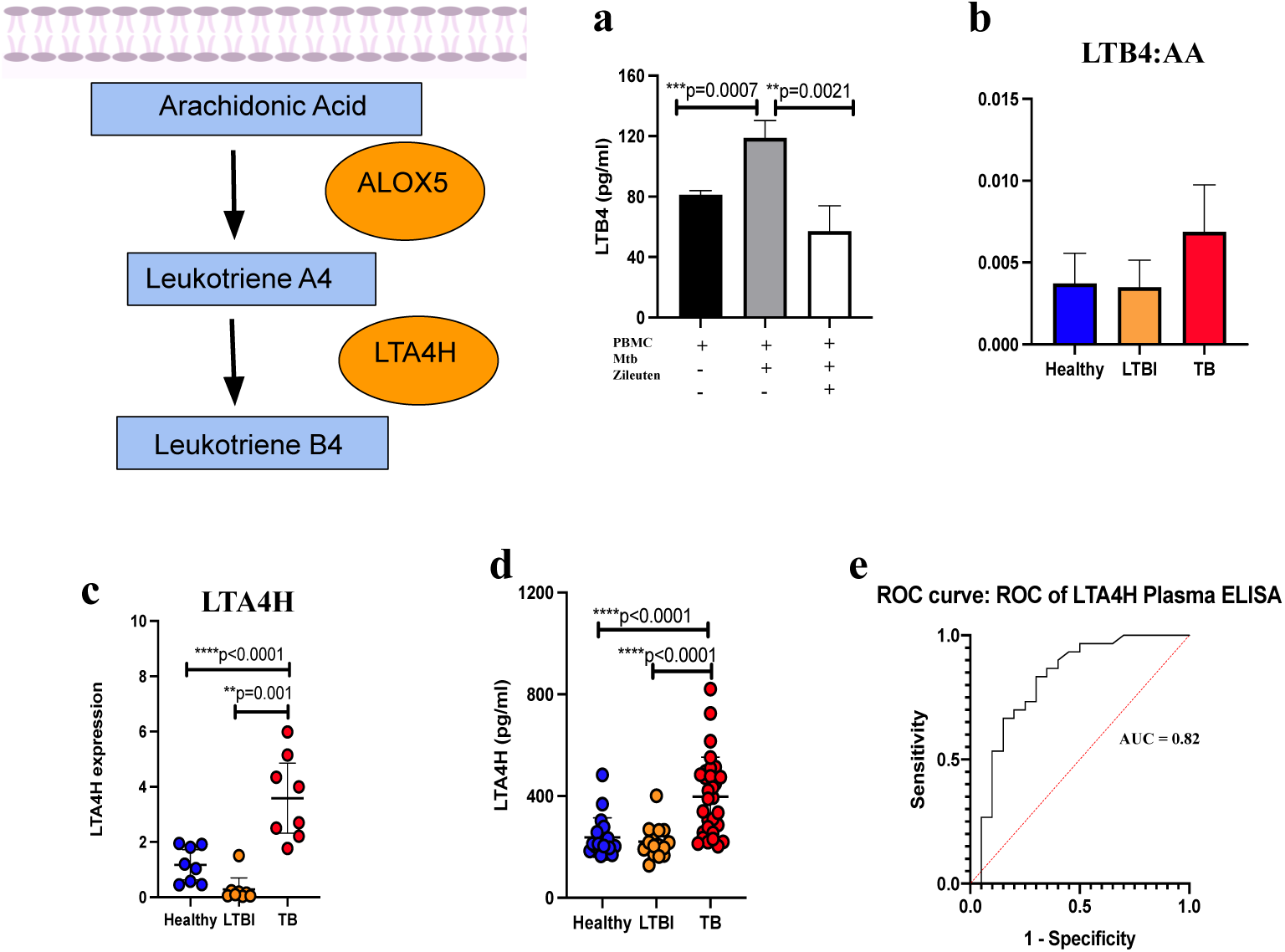
ALOX5 activates LTA4H/LTB4 mediated signaling axis in TB. **a**, LTB4 was quantified using ELISA kit on plasma samples from healthy, latent, and active TB participants. **b**, A ratio of LTB4: AA showing the expression levels of LBT4 in relation to AA consumption was measured using ELISA kit on plasma samples from healthy, latent, and active TB participants. **c,d,** Expression levels of the catalytic enzyme, LTA4H was measured using both qPCR and ELISA from blood samples of healthy, latent, and active TB participants. **e**, The representative ROC curve highlighting the power of LTA4H as a potential predictive biomarker to discriminate between healthy and active TB participants. Error bars denote mean ± SEM. *P < 0.05; **P < 0.01; ***P < 0.001; Student’s t-test was used to measure the significance of the differences between groups. The Area Under Curve (AUC) was used to measure the power of LTA4H in discriminating between groups. The value of 0 indicates a perfectly inaccurate measure and a value of 1 reflects a perfectly accurate measure.

### LTA4H/LTB4 correlate with systemic inflammation in TB infection

To characterize the relationship of LTA4H/LTB4 signaling with TB-induced inflammation, we performed correlation analyses of LTA4H plasma protein with circulatory cytokines/chemokines and growth factors in matching LTBi active TB participants. From the 27 mediators analyzed, pro-inflammatory cytokines/chemokines IP-10, IL-1RA, TNF-α, IL-6, IL-8, and MIP-1b directly correlated with LTA4H (Figures **5a** **-f**). A similar direct correlation was observed between the LTB4 with TNF-α, MIP-1b and IL-8 (figure S4). Interestingly, LTA4H inversely correlated with anti-inflammatory mediators, RANTES and Eotaxin (Figures **5g** and **h**), which also had an inverse relationship to LTB4 (figure S4). Altogether, the data confirm a direct association of LTA4H/LTB4 signaling axis, downstream of ALOX5, with TB induced systemic inflammation.

**Figure 5:**
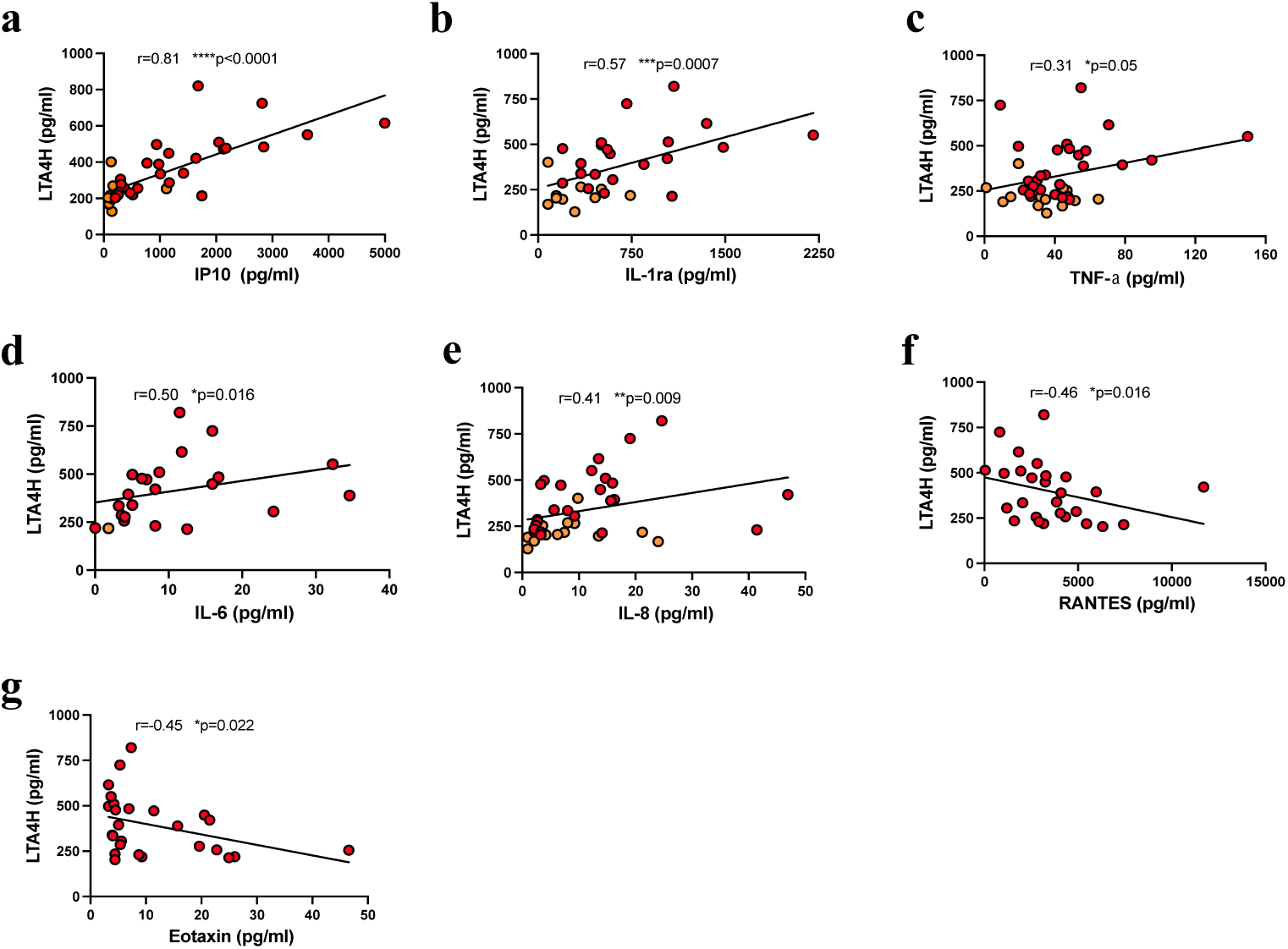
LTA4H/LTB4 correlate with systemic inflammation in TB infection. **a - h,** The expression level of LTA4H measured using ELISA, was correlated with circulatory cytokines/chemokines and growth factors including IP-10, IL-1RA, TNF-α, IL-6, IL-8, MIP-1b, RANTES, and Eotaxin from matching active TB participants. A Pearson correlation coefficient was used to assess the linear relationship between LTA4H and the inflammatory cytokines.

### Pharmaceutical interception of the ALOX5 signaling pathway results in reduced inflammatory responses during *Mtb* infection

Given the association of ALOX5/LTA4H signaling with TB inflammatory responses, we sought to investigate whether this pathway drives inflammation. We utilized Zileuton and SC-57461A, pharmaceutical inhibitors of ALOX5 and LTA4H respectively, to demonstrate the causative role of these proteins on inflammatory responses during *Mtb* infection *in vitro*. Infection of PBMCs with *Mtb* H37Rv resulted in significantly increased pro- and anti-inflammatory cytokines at 24 and 72 hours post infection (Figure **6**). The inhibition of ALOX5 using Zileuton, resulted in a significant reduction of key pro-inflammatory cytokines including TNF-α, IL-1β and IFN-γ, (Figures **6a** **-c**). We observed a similar pattern of cytokine reduction with inhibition of LTA4H using SC-57461A (Figure **6a-c**). The reduction of the inflammatory cytokines due to inhibition of either ALOX5 or LTA4H was concentration dependent. The results confirm that ALOX5/LTA4H pathway drives TB mediated inflammation. Interestingly, the inhibition of both ALOX5 and LTA4H also resulted in reduction of IL-10, a known anti-inflammatory cytokine (Figure **6d**). This suggests that the ALOX5/LTA4H-driven inflammation may be regulated by a feedback mechanism involving IL-10.

**Figure 6:**
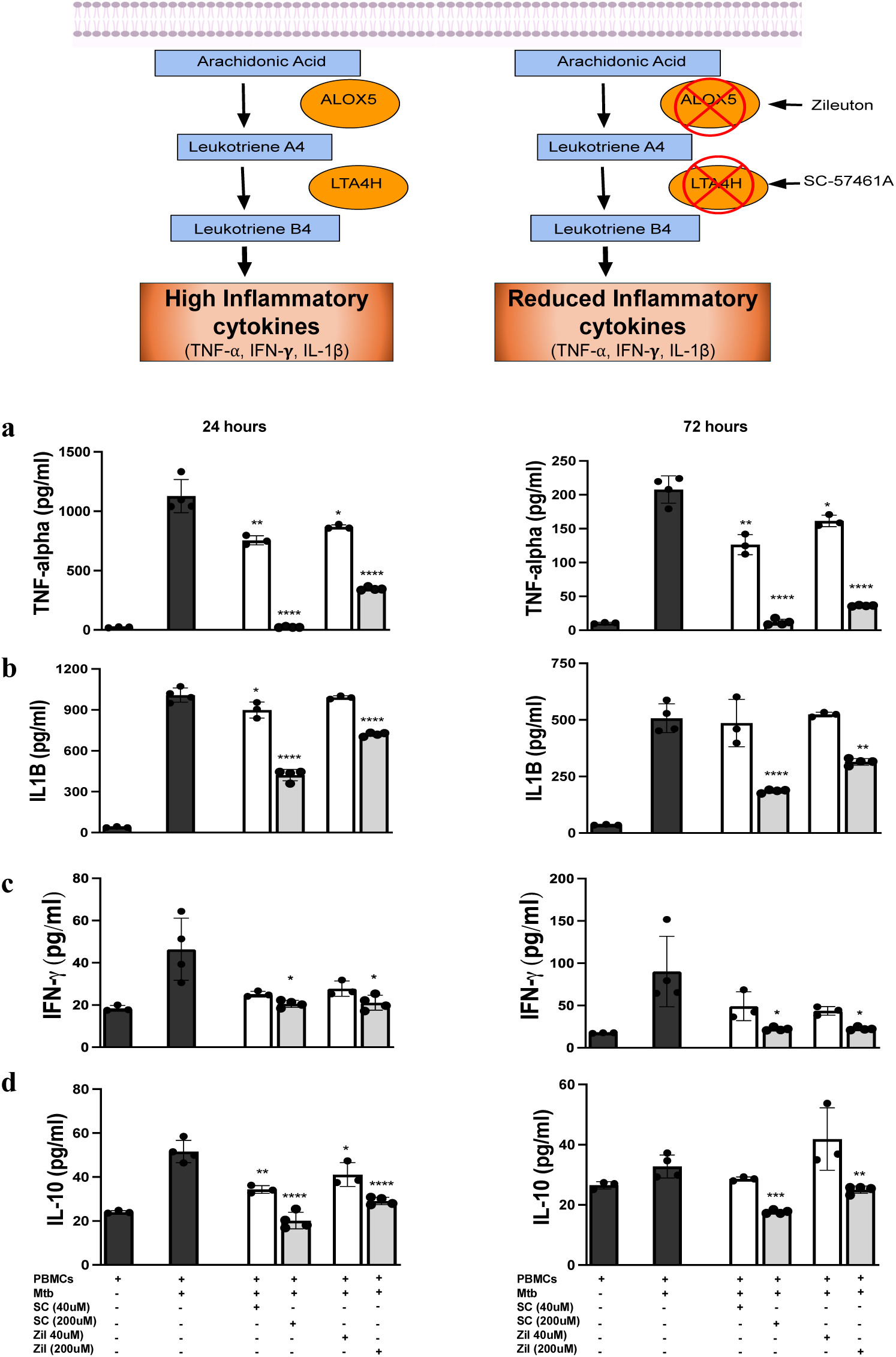
Pharmaceutical interception of ALOX5 signaling pathway results in reduced inflammatory responses during *Mtb* infection. Assessment of ALOX5 signaling on TB induced inflammatory responses using *in vitro* assay. Healthy PBMCs were infected with a virulent strain of H37Rv *Mtb* using MOI of 10 for 24- and 72-hours period (Black bars). **a,b,c,d,** The LTA4H (SC-57461A) and ALOX5 inhibitors (Zileuton) were added at a concentration dependent manner, 40uM (white bars) and 200uM (Grey bars). Inflammatory cytokines including TNF-α, IL1-β, IFN-γ, and IL-10 were measured using ELISA kit. Error bars denote mean ± SEM. *P < 0.05; **P < 0.01; ***P < 0.001; Student’s t-test was used to measure the significance of the differences between groups.

### ALOX5/LTAH4 signaling spatially associates with TNF-α in TB diseased lung

Because ALOX5 and LTA4H operate in the same signaling pathway culminating in release of inflammatory cytokines, we reasoned that these proteins will co-localize within caseous TB granulomas. Using H&E as well as immunofluorescence staining, we found that LTA4H was more abundant in the inflammatory border inside the caseum (Figures **7a and b****; S2d and e**). ALOX5 staining revealed a distribution similar to LTA4H within the same caseous granulomas (Figures **7b and c**). Given our earlier observation on the ALOX5/LTA4H-driven inflammation in plasma and PBMCs, we sought to investigate spatial association of these proteins with TNF-α in the lung tissue. Our staining revealed that TNF-α was uniquely released within the areas of ALOX5/LTA4H enrichment, suggesting that the cytokine is released by ALOX5/LTA4H-expressing cells (Figures **7d and e**). The expression pattens of LTA4H/ALOX5 and TNF-α were confirmed using QuPath software to demonstrate an increased number of cells co-expressing these proteins in the same region of a caseous granuloma (Figure **7f**). Interestingly, we observed co-staining of ALOX5 and TNF-α at individual cell level (Figure **7g****; S5**), confirming that ALOX5-expressing cells are the source of TNF-α-mediated inflammation. These cells were further confirmed to be the inflammatory macrophages as demonstrated with the co-staining of CD68, ALOX5 and TNF-α (Figure **7h**). Our data confirm spatial localization of the ALOX5/LTA4H pathway with inflammation in TB diseased lung.

**Figure 7:**
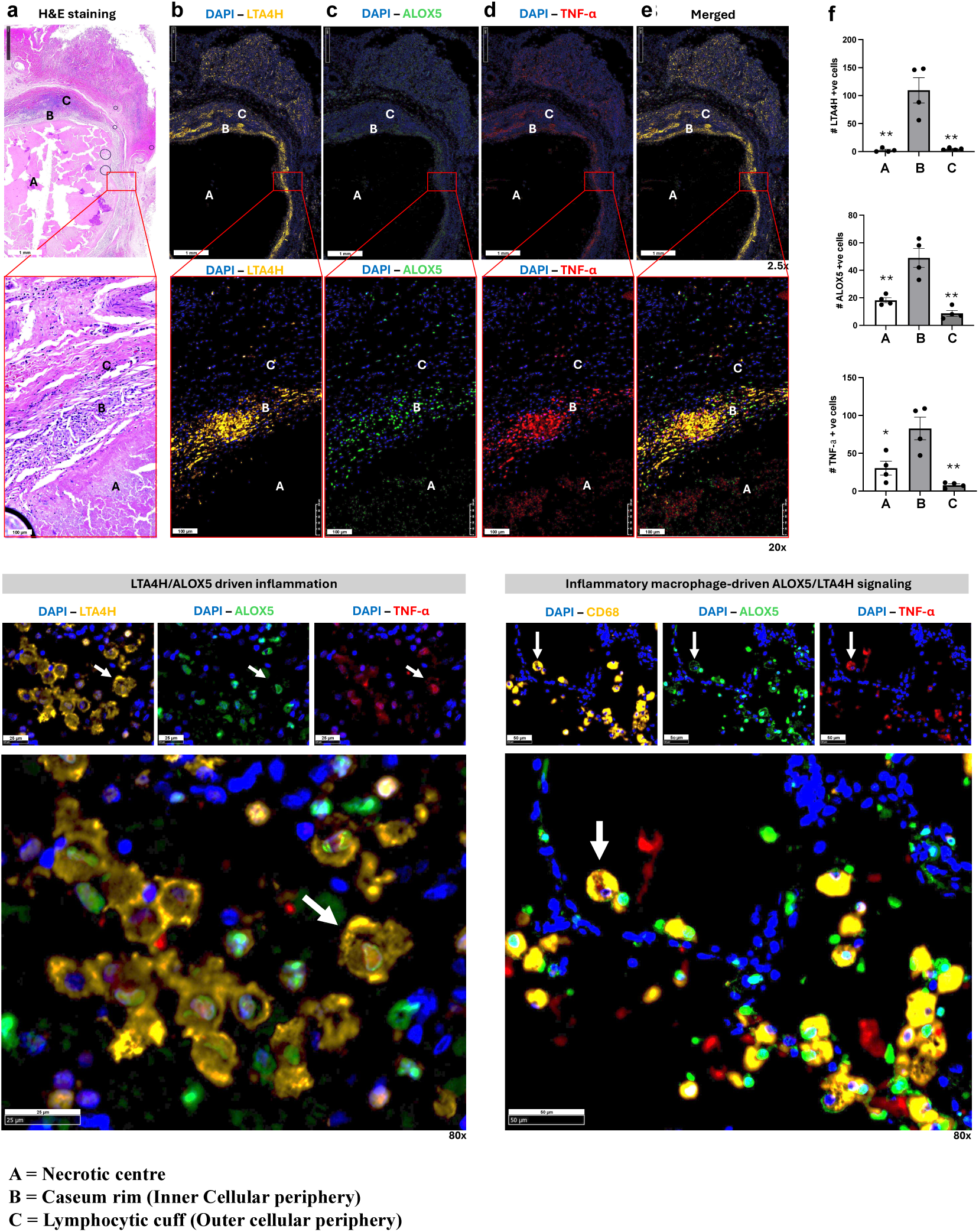
ALOX5/LTAH4 signaling spatially associates with TNF-α on TB diseased lung. **a**, A caseous TB granuloma was identified using H&E staining from a diseased lung tissue with necrotic (A), inflammatory ring (B), and lymphocytic cuff (C) regions outlined. **b,c,d,** Immunofluorescence (IF) staining was used to show spatial expression of LTA4H, ALOX5, and TNF-α in a caseous TB granuloma. **e**, Merged image demonstrating the colocalization of LTA4H, ALOX5, and TNF-α in a caseous TB granuloma from the IF staining. **f**, Quantification LTA4H, ALOX5, and TNF-α expressing units (cells) using QuPath software in the necrotic, inflammatory ring, and lymphocytic cuff of a caseous TB granuloma. **g,** IF staining of LTA4H, ALOX5 and TNF-α to demonstrate their co-localization within the same cell populations. **h,** IF staining of CD68, ALOX5 and TNF-α to demonstrate their co-localization within the same cell populations. Error bars denote mean ± SEM. *P < 0.05; **P < 0.01; ***P < 0.001; Student’s t-test was used to measure the significance of the differences between groups.

### Pharmaceutical inhibition of ALOX5 signaling is a potential therapeutic target against exacerbated granulomatous immunopathology during *Mtb* infection

With the demonstrated role that ALOX5/LTA4H signaling contributes to inflammation associated with caseous TB granulomas, we believed this could be key to controlling exacerbated lung pathology during the disease. To investigate this, C3HeB/FeJ mice were utilized to demonstrate the effect of ALOX5 inhibition on TB immunopathogenesis. The mice were infected with H37Rv *Mtb* strain (500 CFU/mouse) followed by treatment with Zileuton (100mg/kg) from day 5 post infection (Figure **8a**). Analyses at day 56 revealed no observable differences in body (Figure S6) or lung weight between the zileuton treated and the untreated control mice (Figure **8b**). However, in terms of the pulmonary immunopathology, the zileuton treated C3HeB/FeJ mice had significantly reduced granuloma lesion size than the control group (Figure **8c**). Further analysis of lung density maps using QuPath software showed significant reduction of inflammatory cellular infiltrates in the TB-induced lesions of zileuton treated mice compared to untreated mice (Figures **8d** and **e**). This confirms that ALOX5 is a driver of lung pathology in TB. Interestingly, the amelioration of TB-induced immunopathology was accompanied by improved bacterial killing in the zileuton treated mice compared to the control mice (Figure **8f**). Taken together, the data demonstrate that ALOX5 pathway can be targeted to develop host-directed therapy to reduce TB lung damage.

**Figure 8:**
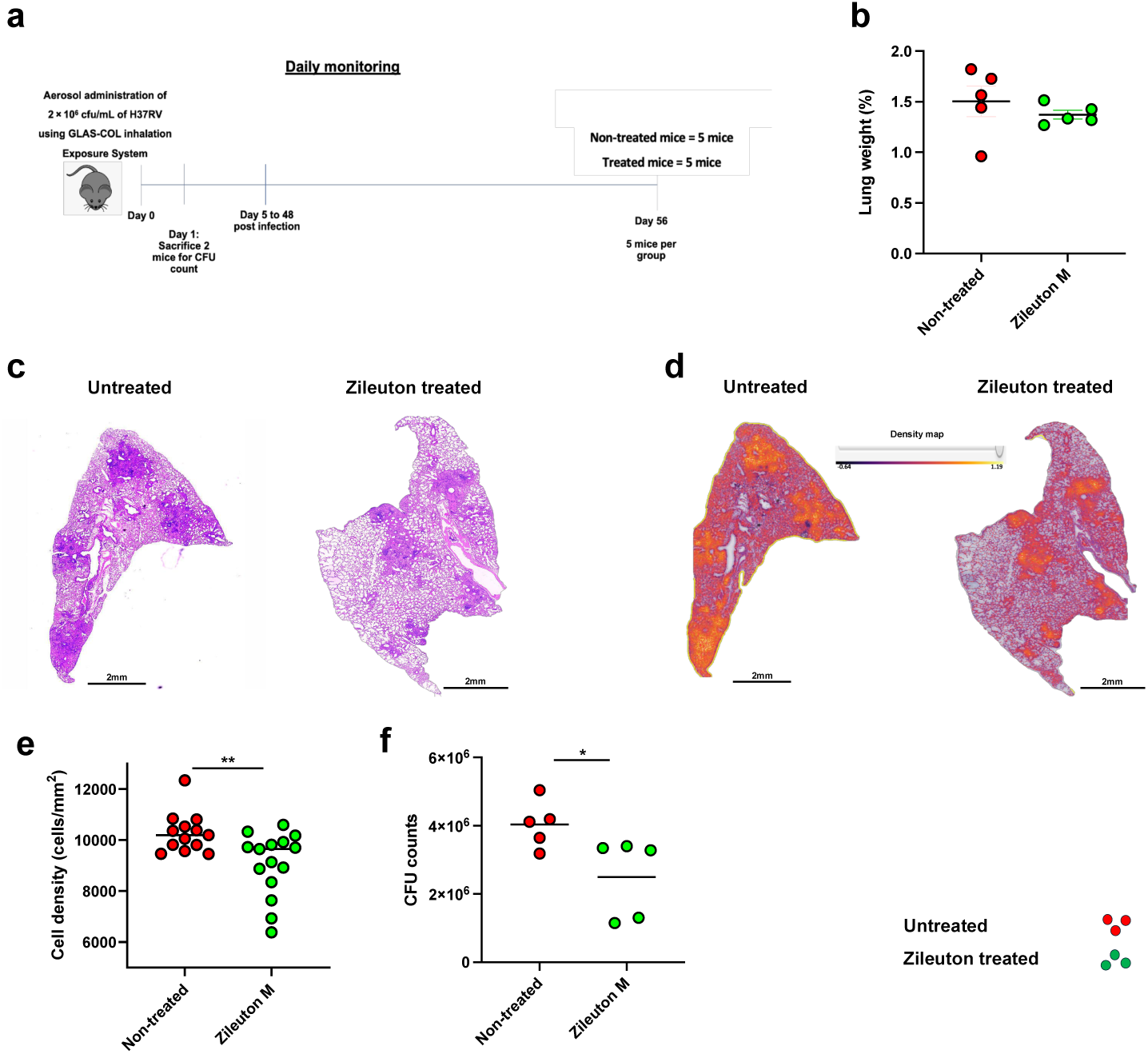
Pharmaceutical inhibition of ALOX5 signaling reduces exacerbated granulomatous immunopathology during *Mtb* infection. To demonstrate the influence of ALOX5 signaling on tissue immunopathology during *Mtb* infection, an *in vivo* model of Kramnik (C3HeB/FeJ) mice was utilized. **a**, A total of 10 Male C3HeB/FeJ mice were infected via inhalation using 2x 10^6^ cfu/ml and 5 of the mice were treated with 100mg/kg of zileuton from day 5 up to day 48 post infection. **b**, Lung tissue weights from treated and non-treated mice were compared at day 56 post infection. **c**, H&E overview of tissue section containing TB-induced granulomatous lesions between treated and non-treated mice on day 56 post infection. **d, e**, Quantification of cellular infiltration around the TB-induced granulomatous lesions between treated and non-treated mice using QuPath software. **f**, Bacterial loads were determined between treated and non-treated mice at day 56 post infection. Error bars denote mean ± SEM. *P < 0.05; **P < 0.01; ***P < 0.001; Student’s t-test was used to measure the significance of the differences between groups.

## Discussion

Tuberculosis remains one of the major global health challenges with those previously exposed presenting persisting pulmonary dysfunction despite being treated for the disease^25^. Furthermore, complications resulting from the disease contribute to a growing global burden of lung dysfunction^33,34^. Our study explores a host factor, ALOX5, which associates with tissue destruction and is demonstrated to have therapeutic potential to treat and/or prevent lung damage during TB.

A unique spatial enrichment of ALOX5, ALOX5AP, and LTA4H proteins was predicted to be highly abundant in severe regions (caseum regions) of TB granulomas^30^. Characterization of lung lesions demonstrated, as previously reported^35^, a variety of TB granulomas that exist in the lung of affected individuals. Using immunopathological assays, ALOX5 was found to be more abundant in severe forms of the TB granulomas (caseous granuloma) compared to solid forms. Furthermore, the host protein was uniquely abundant in the inflammatory layer bordering the caseum centre. Taken together, our data show an association of ALOX5 with inflammatory regions in caseous granulomas, potentially contributing to lung tissue destruction during TB.

Interestingly, the inflammatory layer of caseous TB granulomas was predominantly composed of histiocytes which stained positive for ALOX5. Among the lesions identified, one had increased infiltration of the histiocytes in the necrotic centre, which also stained positive for ALOX5. This led to the hypothesis that, ALOX5 signaling is predominantly mediated by macrophage populations during TB. A unique population of macrophages, which co-stained for both CD68 and ALOX5 was found in the inflammatory layer, confirming the cells as drivers of ALOX5 signaling. Therefore, the data suggested macrophages as the predominant drivers of ALOX5 mediated signaling during TB.

Furthermore, ALOX5 was demonstrated to associate with clinical TB as evidenced by its increased enrichment in active TB patients compared to healthy and latently infected TB participants. The data suggested the host factor as a potential predictor of clinical disease during TB. However, ALOX5 was also demonstrated to be a potential biomarker in low-grade glioma, indicating that it is not TB specific^36^. Therefore, it can be reasoned that the expression of the host factor, primarily associates with tissue remodeling (disease pathogenesis) during disease complications including TB and low-grade glioma.

The host factor, ALOX5, has been shown to be involved in immune- and inflammation related pathways during disease^27–29,36^. Given its association with lung tissue damage, it was also believed to drive inflammatory responses that contribute to tissue pathology. A subset of macrophages expressing ALOX5 was shown to associate with the inflammatory regions in TB diseased lung. Since macrophages are well known for their role in protective inflammatory responses during *Mtb* infection^37^, the ALOX5 was hypothesized to mediate macrophage-driven inflammation via LTA4H during TB. An ALOX5-dependent increase in LTB4 was therefore, shown during *Mtb* infection. This was associated with increased inflammatory cytokines/chemokines in active TB participants. Interestingly, the mediator was observed to inversely correlate with type 2 mediators, RANTES and EOTAXIN, suggesting the role of the host factor during type 1 inflammation. Several studies have also highlighted the significance of ALOX5 signaling on inflammatory responses during *Mtb* infection. In patients suffering from TB meningitis, a single nucleotide polymorphism in the LTA4H locus was associated with increased inflammatory cells, and an improved response to adjunct anti-inflammatory therapy^38,39^. Furthermore, a previous study also demonstrated an improved protection against *Mtb* infection in the absence of ALOX5 signaling using murine models^40^. These findings, therefore, encouraged the notion of ALOX5 signaling, via LTA4H, as a potential contributor to pathogenic inflammatory responses during TB.

The role of ALOX5 signaling to inflammatory responses during TB, was confirmed using *in vitro* assays. The disruption of the signaling pathway, through ALOX5 and/or LTA4H inhibitors, contributed to a significant reduction in pro-inflammatory responses. This was consistent with previous studies, which also showed, using murine models, that ALOX5 signaling does contribute to pro-inflammatory responses during *Mtb* infection^40,41^. Although the increase in ALOX5 signaling was hypothesized to also associate with reduced anti-inflammatory cytokines. This was not the case as evidenced by an increase in IL-10 responses during *Mtb* infection. Furthermore, the response was reduced by the disruption of ALOX5/LTA4H signaling. Interleukin 10 is a pleiotropic cytokine, which can inhibit macrophage activation leading to inhibition of Th1 cell development, proliferation and cytokine production^42–44^. Therefore, the increase in IL-10 production during the infection was a result of a feedback mechanism mediated to encounter exacerbated inflammation during TB. Given the role of ALOX5 signaling in inflammatory responses, the pathway was suggested, using *in vivo* models, to drive increased lung inflammation and tissue damage during *Mtb* infection^40,45^. The current study found that subsets of macrophages expressing ALOX5 in the inflammatory layer of caseous granulomas, were also expressing TNF-α in TB patients. This confirmed the role of macrophages, through ALOX5 mediated signaling, in driving increased lung inflammation associated with lung pathology in TB patients.

To understand the role of ALOX5 mediated inflammation in tissue pathology, Kramnik (C3HeB/FeJ) mice were utilized, since they can develop TB-induced necrotic granulomas when infected with a pathogenic strain of *Mtb*^46^. The disruption of ALOX5 signaling using zileuton, resulted in reduced lung lesions in *Mtb* infected mice. This demonstrated a role for the signaling pathway to contribute to the development of lung lesions during *Mtb* infections. Formation of the lesions was attributed to increased caseous necrosis of macrophages during TB. This is because, it was shown previously, that excess LTA4H, results in increased TNF-α levels which in turn cause programmed necrosis of macrophages in zebrafish^38,47^. Furthermore, continued macrophage necrosis has been demonstrated to contribute to the development of caseous granulomas during TB^12,13,48,49^. Therefore, taken together, the work demonstrates a role for macrophage driven inflammation via ALOX5/LTA4H signaling, in contributing to TB-induced lung pathology. The regulated lung pathology culminated in improved bacterial killing as evidenced from the Kramnik mouse model utilized.

We, therefore, present a case to target mediators of damage such as ALOX5, in developing HDTs to manage TB induced lung damage. This will provide an adjunct drug candidate with first line drugs (i.e., rifampicin) to help reduce the treatment period, as well as improve the long-lasting effects commonly seen in individuals who have been cured from TB disease.

## Methods

### Study setting and collection of blood and lung samples

Lung samples were obtained from thirty-nine participants (n = 39) presenting with a range of TB associated lung pathological complications, at a collaborating hospital called King Dinizulu Hospital Complex based in Durban, South Africa. Demographics and clinical characteristics of the study participants are listed in Supplemental Table 1. We confirm that all research was performed in accordance with relevant guidelines/regulations. Informed consent was obtained from all participants and/or their legal guardians. The study was carried out at Africa Health Research Institute, under an approved lung study program through Biomedical Research Ethics Committee (BREC) at the University of KwaZulu-Natal (BE019/13). Each consenting participant was allocated a personal identifying data (PID). Lung tissue biopsies ranging from non-affected (no pathological damage) to mild and severely damaged, were carefully isolated and preserved in 4% formaldehyde solution.

For blood sample study, we recruited participants from the KwaDabeka clinic and Prince Cyril Zulu Communicable Disease Centre, in the eThekwini district of Durban in KwaZulu-Natal (KZN), South Africa. This study was approved by the Biomedical Research Ethics Committee (BREC) at the University of KwaZulu-Natal (BE022/13/ BE0000365/2021). We confirm that all research was performed in accordance with relevant guidelines/regulations. Informed consent was obtained from all participants and/or their legal guardians. We recruited TB patients who were newly diagnosed as GeneXpert positive (n=30), and LTBI (n=20) who were QuantiFERON (QFT) positive and healthy individuals (n=20) (QFT negative). All participants were treatment naïve at the time of sample collection. Clinical characteristics of all 70 participants used in this study are listed in Supplemental Table 2. Whole blood samples were collected and processed as previously described^34^. Lung samples were obtained from individuals undergoing elective pneumonectomy at King Dinizulu Hospital Complex and were collected through an ongoing study at the Africa Health Research Institute (BE019/13).

### Sandwich ELISA

The ELISA assay was used to quantify levels of cytokine responses according to the manufacturer’s instructions from plasma and cell culture supernatant. BD OptEIA ELISA kits (BD Biosciences, USA) were used to measure cytokine levels including IL-1β, TNF-α, IL-6, IL-10, MCP, IFN-γ) according to the manufacturer’s instructions as outlined previously^51^. The LTA4H was measured using Human LTA4H ELISA Kit (ThermoFisher Scientific, cat. no.: EH308RB), and the LTB4 was measured using Human LTB4 ELISA Kit.

### Colorimetric assay

Colorimetric assay was used to quantify the levels of AA, ALOX5, LTA4H in plasma and supernatant from cultured PBMC samples according to the manufacturer’s instructions. The following kits were used, Human Arachidonic Acid ELISA Kit (Novus Biologicals, cat. no.: NBP2-59872). Human 5-Lipoxygenase ELISA Kit (ThermoFisher Scientific, cat. no.: NBP2-66367). The assay was based on a competitive binding between target protein (AA, ALOX5) and an acetylcholinesterase (AChE) conjugate (tracer) for binding to a monoclonal antibody specific to the protein of interest. The antibody-protein complex was then bound to goat polyclonal anti-mouse/rabbit IgG that was attached to the wells from the plates provided. Any unbound reagents were washed off and an Ellman’s Reagent containing AChE was added. The enzymatic reaction resulting in a distinct yellow colour, was measured using an OD reader with a wavelength of 414nm. The intensity of the yellow colour was proportional to the amount of tracer bound to the well and was inversely proportional to the amount of target protein present in the test sample. The following formula was used to quantify the amount of target protein detected using the Kit: % Cross reactivity = [(50% B/B_0_ value for the primary analyte)/ 50% B/B_0_ value for the potential cross reactant)] x 100%.

### Multiplex ELISA

The assay was performed as previously described^52^ using Bio-Plex 27 human cytokine panel kit (Hercules, CA, USA), on plasma samples obtained from healthy, latently infected, and active TB participants. A Bio-Plex 200 plate reader (CA, USA) was used to measure the 27 cytokine and chemokine including, interleukin (IL)-1β, IL-1 receptor agonist (RA), IL-2, IL-4, IL-5, IL-6, IL-7, IL-8, IL-9, IL-10, IL-12, IL-13, IL-15, IL-17, interferon-γ (IFN-γ), tumour necrosis factor (TNF-α), Eotaxin, RANTES, granulocyte macrophage colony stimulating factor (GM-CSF), granulocyte colony stimulating factor (G-CSF), fibroblast growth factor (FGF) basic, interferon gamma inducible protein (IP-10), monocyte chemoattractant protein-1 (MCP-1), platelet derived growth factor (PDGF-bb), macrophage inflammatory protein alpha (MIP-1α), MIP-1β, and vascular endothelial growth factor (VEGF). The assay included duplicates of both the plasma and standard samples. A standard curve was generated using Bio-Plex-manager software version 6, and this was used to work out the cytokine/ chemokine concentration levels in the plasma samples.

### Histology

#### Tissue processing

Lung tissue samples collected were processed using Microm STP 120 Spin Tissue Processor (ThermoFisher Scientific, USA) in accordance with the manufacturer’s instructions to remove 10% neutral buffered Formalin solution (ref. no. HT501128-4L; SIGMA-Aldrich, USA) that was used to store the samples. The tissue samples were then embedded into wax blocks using HistoStar Embedding Station. The samples were then cut into 5μM sections and embedded on to the X-tra^TM^ Adhesive micro slides (ref. no.3800200AE; Leica Biosystems, USA).

Prior to tissue staining, paraffin solution was removed from the embedded tissue sections using serial wash steps in the following order: Incubation of slides for 2x 5mintes in Xylene, then dehydration in 2x 5 minutes of 100% followed by 95% and 70% of ethanol solution. The slides were then incubated in distilled water for 5 minutes.

#### For H&E staining

the slides were subjected to hematoxylin staining for 5 minutes, then washed in running tap water for 5 minutes. The sections were then stained with eosin for 2 minutes and then rinsed in 95 % ethanol for 30 seconds, followed by 100 ethanol for 1 minute. To mount the slides with a cover slip, the slides were rinsed in Xylene for 1 minute and then a cover slip was added using mounting media.

### Immunohistochemistry (IHC) staining

The tissue samples embedded on slides were subjected to deparaffinization as above, and antigen retrieval was performed by running the samples with 1L of EnVision™ FLEX target retrieval solution high pH (30ml of concentrate diluted in 970ml of distilled/ DI water) using PT link. Following this, the slides were blocked using peroxidase block (S202386-2, Agilent, USA) for 5 minutes, and then washed 2 times for 5 minutes (2x 5minutes) using TBS buffer. The tissue samples were blocked again using protein block (0.05g BSA + 1% goat serum in TBS wash buffer). The slides were washed as above (2x 5 minutes), and a primary antibody of interest was added and incubated for 45 minutes at room temperature.

Tissue sample staining: For blocking, 1% goat serum was added to the tissue sections for 20 minutes to block non-specific immunostaining. The primary antibody of interest was added in an appropriate concentration that is recommended by the manufacturer or previous literature and then incubated for 1 hour at room temperature (RT). The antibody was washed from the slides using Tris buffer and the sections soaked in Tris buffer for 10 minutes (2x 5 min washes). The slides were then incubated with corresponding biotinylated goat anti-species antibody for 10 minutes, followed by rinsing the slides using Tris buffer for 10 minutes (2x 5 minutes washes). The detection enzyme solution (streptavidin peroxidase) was added, and the slides incubated in a dark humid chamber for 5 minutes. The slides were rinsed in Tris buffer for 10 minutes (2x 5 minutes washes). A solution of chromogen, 3,3′-diaminobenzidine (DAB) at a concentration of 1mg/ml in Tris buffer with 0.016% fresh hydrogen peroxide was added to the slides and incubated for 8 minutes. The DAB was washed from the slides using tap water for 1 minute.

Addition of counter staining: The slides were dipped into a solution of hematoxylin (or Methyl Green) that is diluted 1:1 in distilled water, and then incubated for 1 minute to produce a very light nuclear counterstain. The slides were washed using double distilled water, and then dehydrated by dipping 95% ethanol for 1 minute, then 100% ethanol for 1 minute. The slides were washed 3 times using xylene and a coverslip mounted on the slides. The slides were scanned using Hamamatsu NanoZoomer 2.0RS scanner (ThermoFisher Scientific, USA) to identify the staining of interest.

### Immunofluorescence (IF) staining

For immunofluorescence staining, the embedded tissue sections were subjected to paraffin removal and antigen retrieval using PT Link system. The samples were then washed in 1x EnVision TM Flex wash buffer (DM831, ref.: K8000/K8002, Agilent, USA) for 10 minutes (2x 5 minutes washes). To allow ease of staining, a hydrophobic ped was used to circle around the tissue of interest. The tissue sections were then blocked using endogenous peroxidase blocking buffer (S202386-2, Agilent, USA) for 10 minutes (2x 5 minutes washes) at room temperature (RT) and then washed using the wash buffer for 10 minutes (2x 5minutes). A second blocking solution (0.05g BSA + 0.5ml Goat serum + 4.5ml wash buffer) was added to the sections for 20 minutes and then washed as above. A primary antibody of interest was then to the slides and incubated for 40 minutes at room temperature. Following the incubation, the slides were then washed as above (2x 5 minutes washes). A secondary antibody was conjugated to the primary by adding Opal Polymer (HRP: Ms + Rb) to the tissue and incubated for 20 minutes at room temperature. To label the 1st primary antibody a fluorescent dye of wavelength 494/525nm (OPAL 520 reagent, AKOYA Biosciences, USA) was added at a dilution of 1:200 using 1x Plus Amplification buffer and incubated for 10 minutes. A second antibody of interest was added to the tissue by first exposing the antigens on the tissue using TSC solution (10% of Antigen Retrieval 6 buffer (K800421-2, Agilent, USA) diluted in distilled water. The solution was initially boiled using microwave and the slide were added to solution and microwave heated at 3 different conditions. The first condition was heating the slides for 2 minutes with the microwave set on high, then heated for 5 minutes with the microwave set on medium, and lastly, the slides were heated for 10 minutes with the microwave set on low. Once done, the bowl containing the TSC solution with the slides was placed in running tap water for 20 minutes to cool the solution. To equilibrate the tissue, the slides were rinsed in 1x EnVision TM Flex wash buffer (DM831, ref.: K8000/K8002, Agilent, USA) for 5 minutes at RT. The second primary antibody was then added following the steps as with the first primary antibody, followed by secondary Opal Polymer (HRP: Ms + Rb) antibody, and a second fluorochrome of wavelength 550/570nm (OPAL 570 reagent, AKOYA Biosciences, USA). Using similar steps above, the third primary antibody with a fluorochrome of wavelength 676/694nm (OPAL 570 reagent, AKOYA Biosciences, USA) were added to the slides. To complete the staining, the tissue samples were then stained with DAPI for 5 minutes at room temperature. Once done, the slides were then washed using wash buffer as before and then covered with coverslips using Dako fluorescent mounting media. The slides were scanned and viewed using Hamamatsu NanoZoomer S60, Japan). Antibodies used in this study are listed in Table 2.

**Table 2:** Antibodies used in this study.

| Antibody | Catalog no. | Company | Host | IHC dilution | IF dilution |
| --- | --- | --- | --- | --- | --- |
| Anti-ALOX5 | ab169755 | Abcam | Rabbit monoclonal | 1/100 | 1/100 |
| Anti-COX1 | ab111855 | Abcam | Rabbit polyclonal | - | 1/100 |
| Anti-LTA4H | ab133512 | Abcam | Rabbit monoclonal | - | 1/250 |
| Anti-TNFα, | ab225576 | Abcam | Rabbit monoclonal | - | 1/100 |
| Anti-CD68 | ab213363 | Abcam | Rabbit monoclonal | - | 1/100 |

### RNA isolation and cDNA synthesis

For RNA isolation and cDNA synthesis, 2.5 ml of whole blood was collected in PAXgene™ tubes and RNA was isolated using the Paxgene™ kit (PreAnalytix, Hombrechtikon, Switzerland). Complimentary DNA (cDNA) was synthesized from isolated RNA as per the manufacturer’s instructions (Bio-Rad, Hercules, CA). Briefly, RNA was adjusted to a concentration of 500 ng for cDNA synthesis and an appropriate volume of the reaction mix containing 5-20 µl of nuclease free water, 1µl of RNA and 4µl of iScript 5× was added to each sample for a total reaction volume of 20µl. The T100 thermocycler from Bio-Rad (Hercules, CA, USA) was set to 5 min at 25°C, 30min at 42°C, 5 min at 85°C.

### Real time quantitative polymerase chain reaction for gene expression analysis

Real time quantitative polymerase chain reaction (RT-qPCR) was done to determine the expression of neutrophil specific genes as per the manufacturer’s instruction pertaining to the iTaq™ Universal SYBR green supermix (Bio-Rad, Hercules, CA, USA). The total reaction volume was 10 µl. Briefly, 1µl of cDNA was added to each well with 9 µl of mastermix consisting of the relevant forward and reverse primer at 0,5 µl each, 5 µl of iTaq™ Universal SYBR green supermix (Bio-Rad, Hercules, CA, USA) and 3 µl of nuclease free water. The CFX 96 thermocycler (Bio-Rad, Hercules, CA) was set to the following protocol: 30 Sec at 95°C, 5 sec at 95°C, 30 sec at 56°C for 39 cycles. The melt curve analysis was done at 65-95°C with 0,5 °C increments. Glyceraldehyde 3-phosphate dehydrogenase (GAPDH) was used as the housekeeping gene and all qPCR data was normalized to GAPDH expression.

### Primer design

Primers were designed using the IDT primer design tool, PrimerQuest Tool and sequences were blasted using the BlastN tool on NCBI. Primers were designed for ALOX5, ALOX5AP, LTA4H, COX1 and COX2 (see Table 3).

**Table 3:**
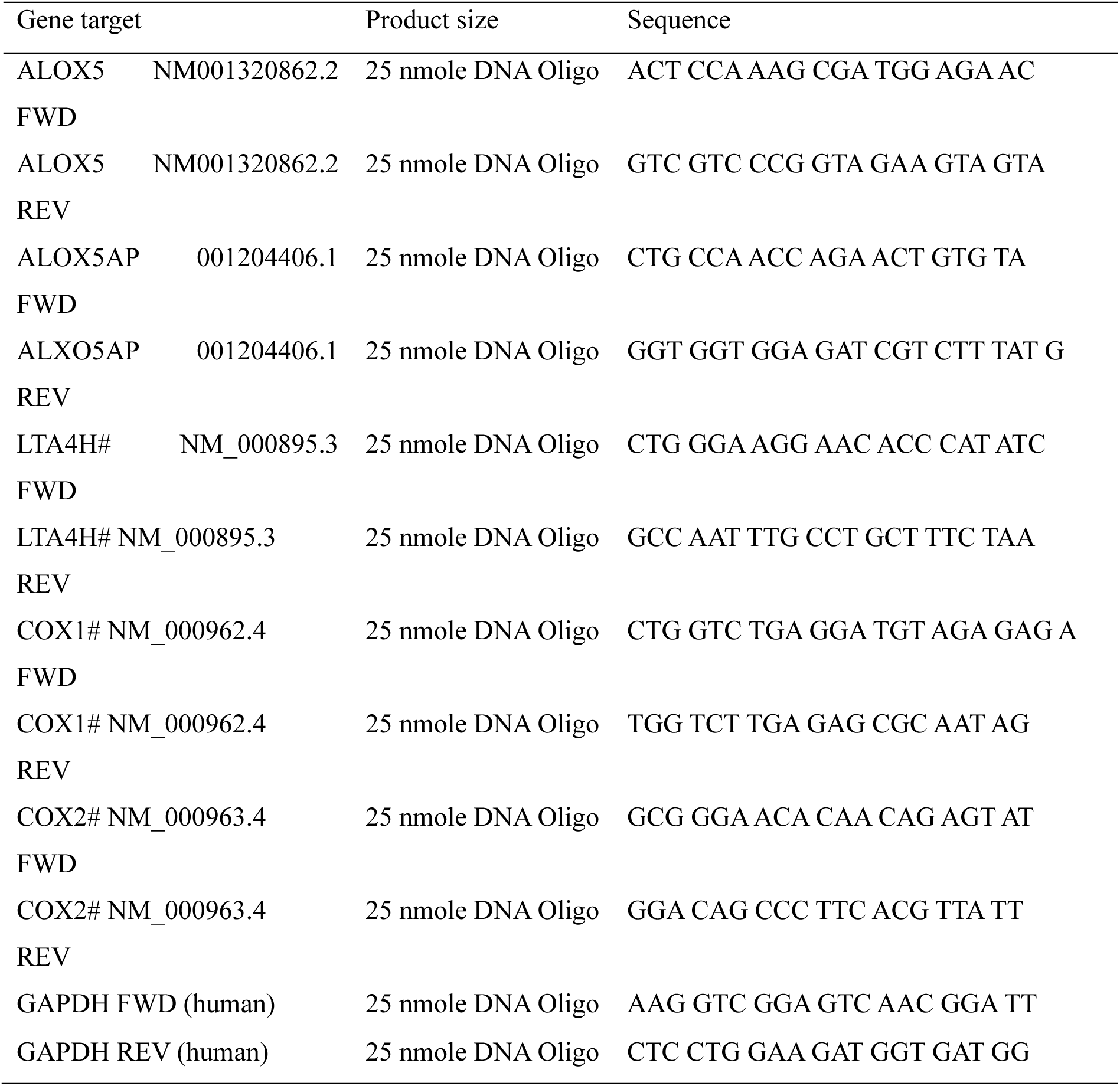
qPCR primer inventory from IDT/WhiteSci.

### Mice

The C3HeB/FeJ mice were donated by the Parihar Lab from University of Cape Town and were originally purchased from the Jackson Laboratories (USA). The animals were bred and housed in Africa Health Research Institute (AHRI) animal facility in accordance with South African National Standard (SANS 10386: 2021. Edition 2) and University of KwaZulu Natal laboratory animal procedures under protocol reference number AREC/00004354/2022.

### Experimental mouse infection

The C3HeB/FeJ mice were infected with a dose of 2x10^6^ H37Rv *Mtb* bacilli using GLAS-COL inhalation system. One day post infection, 2 mice were sacrificed to confirm the bacterial dose of 200 CFU/mouse was delivered. The mice were then treated with 100mg/kg Zileuton (ALOX5 inhibitor) in drinking water from 5 to 48 days post infection.

The CFU counts were confirmed in lung tissue samples harvested from the euthanized mice, following homogenization using 0.01% Tween-PBS. Serial dilutions (5-, 125-, 625-, 3125-fold) were plated on 7H10 agar plates to allow the growth of *Mtb* colonies to confirm the success of the infection and the required dose of CFU/mouse administered.

## Acknowledgements

The study was made possible and carried out successfully due to everyone who took part in it. This includes study participants who consented to the use of clinical samples for research; Operating personnel including surgical doctors and nurses, from King Dinizulu Hospital Complex; AHRI grant office and clinical core personnel who assisted with patient recruitment and sample collection; AHRI staff members who assisted with processing of clinical samples (blood and lung samples); AHRI staff from the Optic and Imaging Core. Dr. Omolara Baiyegunhi, Dr. Duran Ramsuran, Mr Delon Naicker, who provided training and induction to the imaging facility and equipments. Professor Eric J. Rubin for useful discussions and providing useful guidance.

## Author contributions

**Conceptualization:** M.J.M., T.M. **Investigation:** T.M., K.R., D.M., H.N., K.N., T.K.J.L.,

W.S., K.F., K.T., K.L., P.M., A.R., M.J.M. **Methodology:** T.M., K.R., D.M., H.N., K.N.,

T.K.J.L., W.S., K.F., K.T., K.L., P.M., A.R., M.J.M. **Data validation:** T.M., K.R., D.M., M.J.M.

**Original draft:** T.M and MJM. **Review & editing:** T.M., K.R., D.M., K.N., T.K.J.L., W.S., K.F., K.T., K.L., P.M., A.R., R.M., S.P., T.N., S.G., H.N., A.S., M.J.M. **Supervision:** T.M.,

S.G., H.N., A.S., M.J.M. **Resources:** A.S., H.N., S.P., M.J.M. **Funding acquisition:** M.J.M.

## Funding

The study was carried out with the help of funding initiatives awarded to M.J.M. by Wellcome Trust (grant# 206751/A/17/Z); Grand Challenges, an initiative of the Bill & Melinda Gates Foundation (grant #OPP1210776, grant# INV-016239); South African Medical Research Council through its Division of Research Capacity Development under the Mid-Career Scientist Programme received from the South African National Treasury. TM was awarded National Research Foundation Postdoctoral Fellowship.

## Ethics statement

This study was approved by the Biomedical Research and Ethics Committee at the University of KwaZulu-Natal. The blood (BE022/13/BE0000365/2021) and lung tissue (BE019/13) samples were obtained through an ongoing study at the Africa Health Research Institute.

## Conflict of interest

The authors declare no conflict of interests.

---

**Figure S1.**
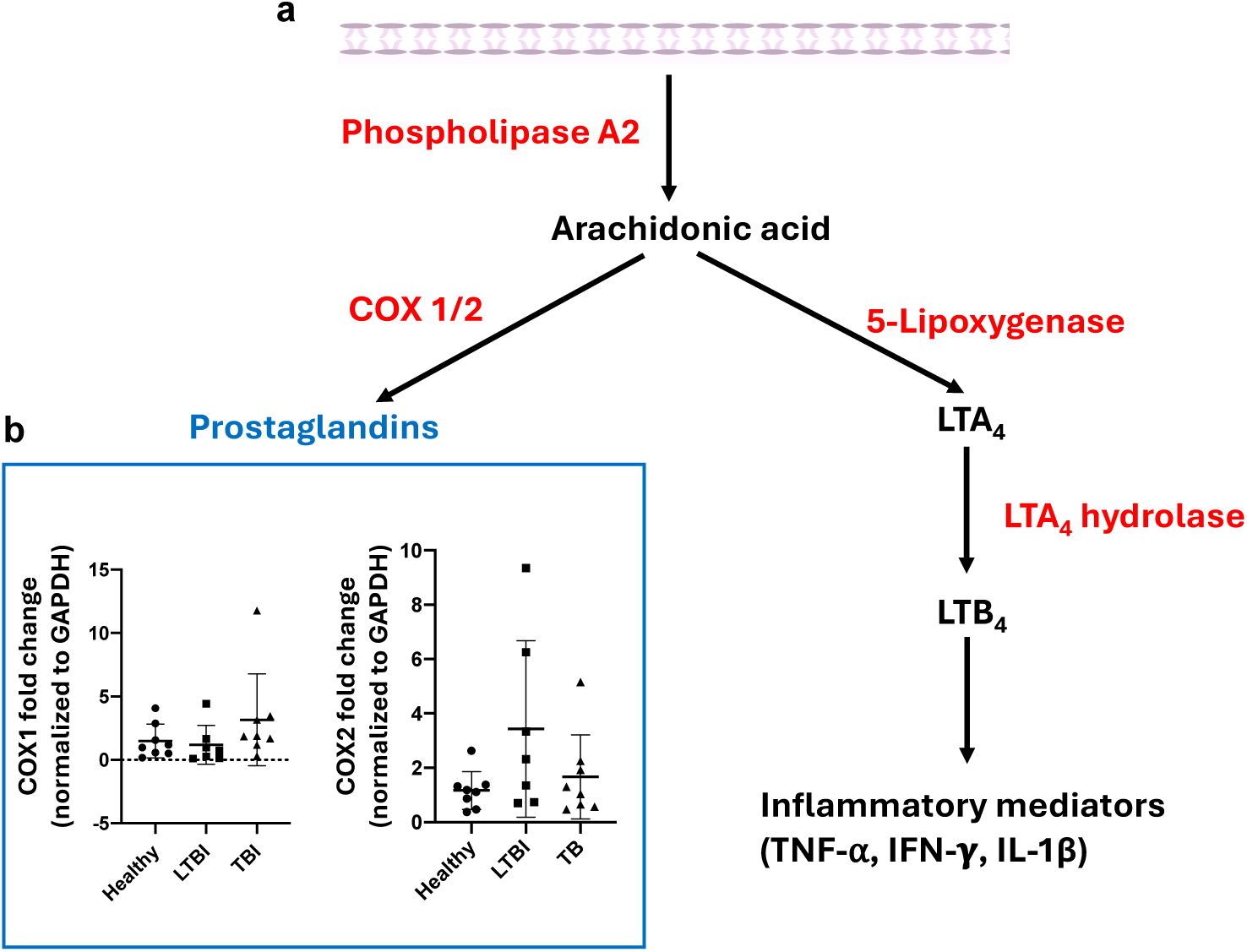
**a**. A schematic flow diagram of arachidonic acid metabolism leading to either 5-lipoxygenase or COX1/2 mediated signaling pathway. b. Expression levels of the catalytic enzymes, COX1 and 2, were measured using both qPCR from blood samples of healthy, latent, and active TB participants.

**Figure S2.**
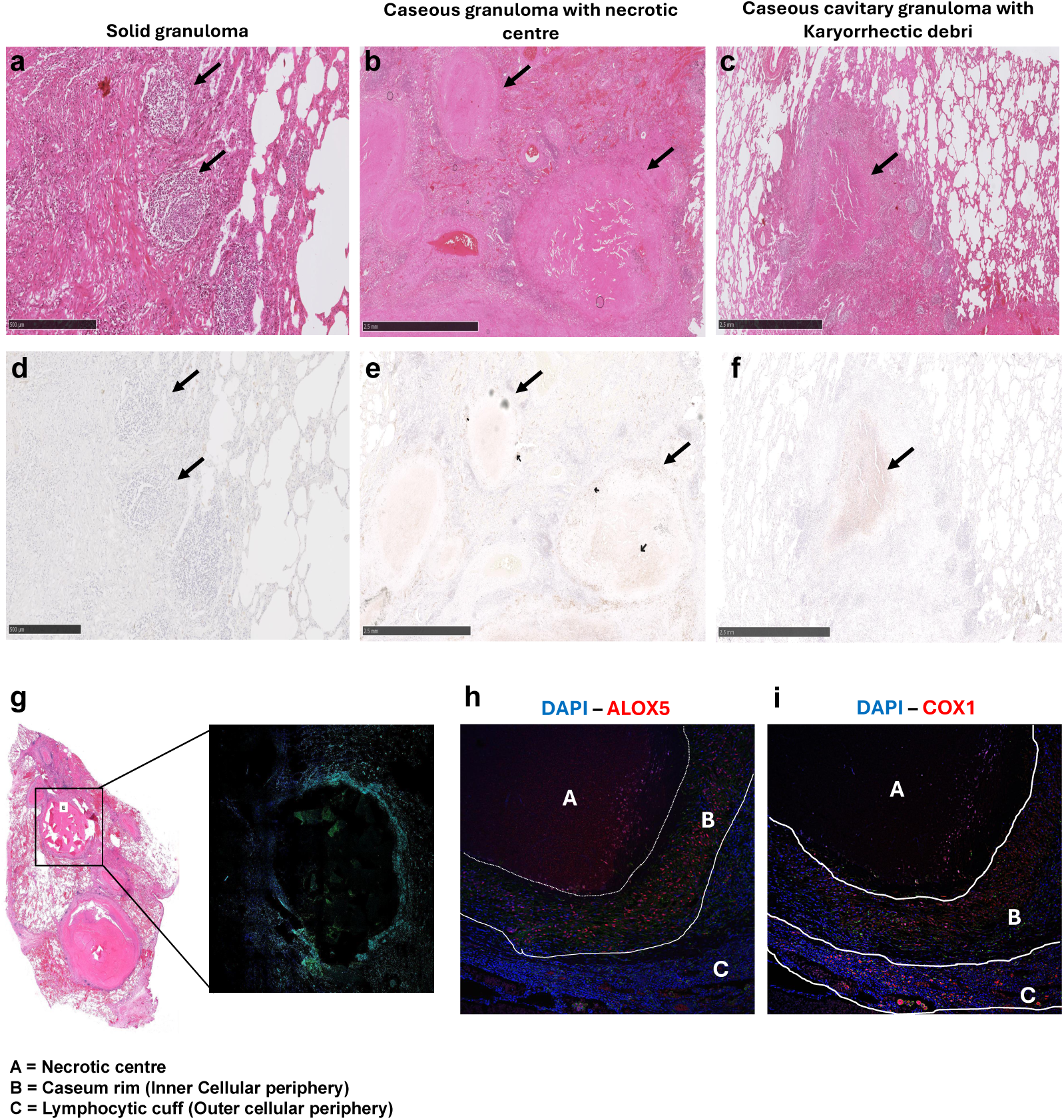
**a** – **c**, H&E overview of tissue section containing varying forms of TB-induced granulomas signifying the degree of damage, from solid to caseous necrotic, caseous cavitary granuloma. **d – f**, Abundance of ALOX5 protein quantified using immunohistochemistry on tissue sections containing varying forms of TB granulomas. **g** and **h**, Spatial expression of ALOX5 on caseous TB granuloma using immunofluorescence (IF) staining. **i**, Spatial expression of COX1 on caseous TB granuloma using immunofluorescence (IF) staining.

**Figure S3.**
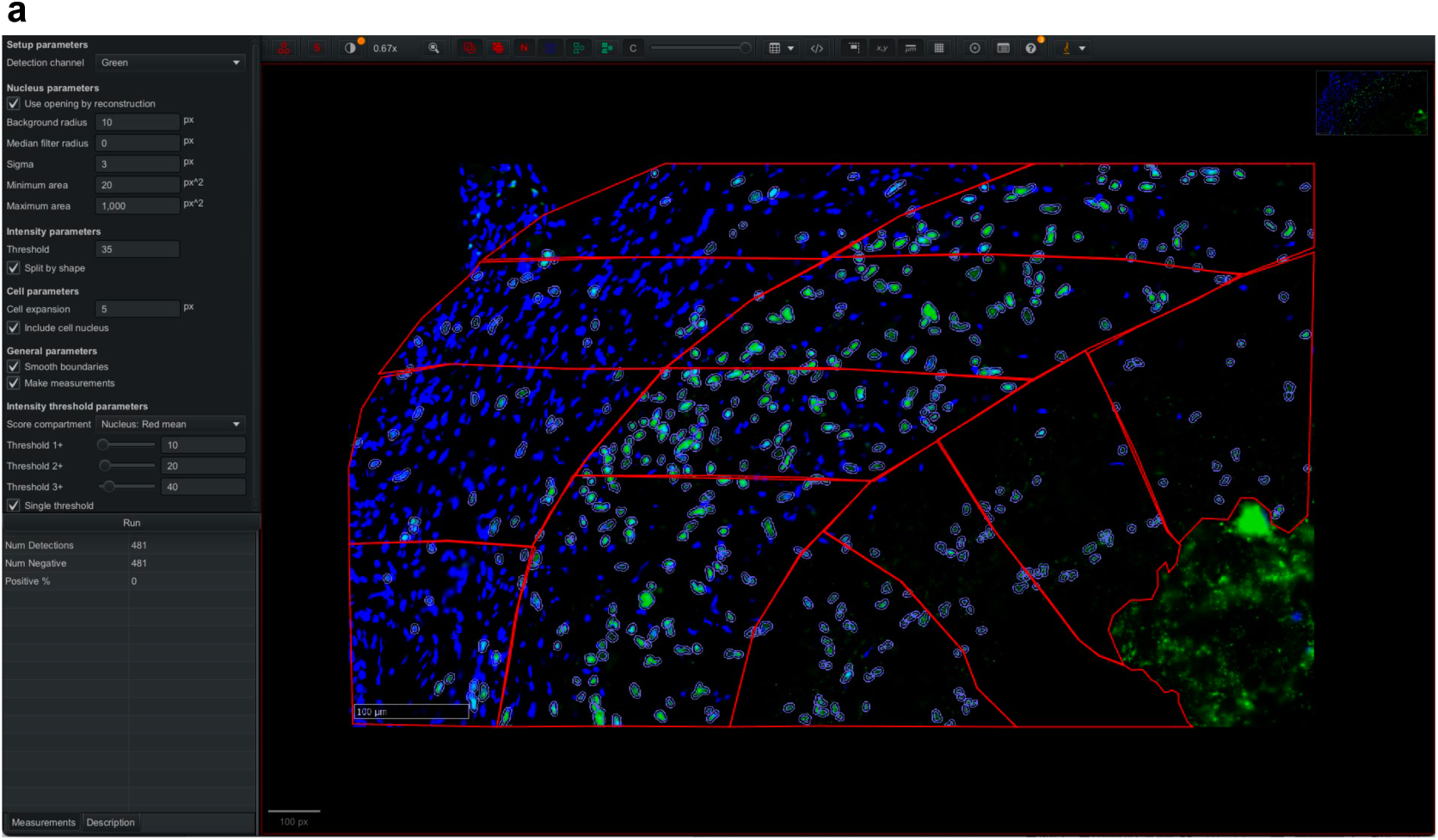
An overview of QuPath software showing caseous granuloma layers sub-divided into 4 sections for quantifications of a protein of interest per layer.

**Figure S4.**
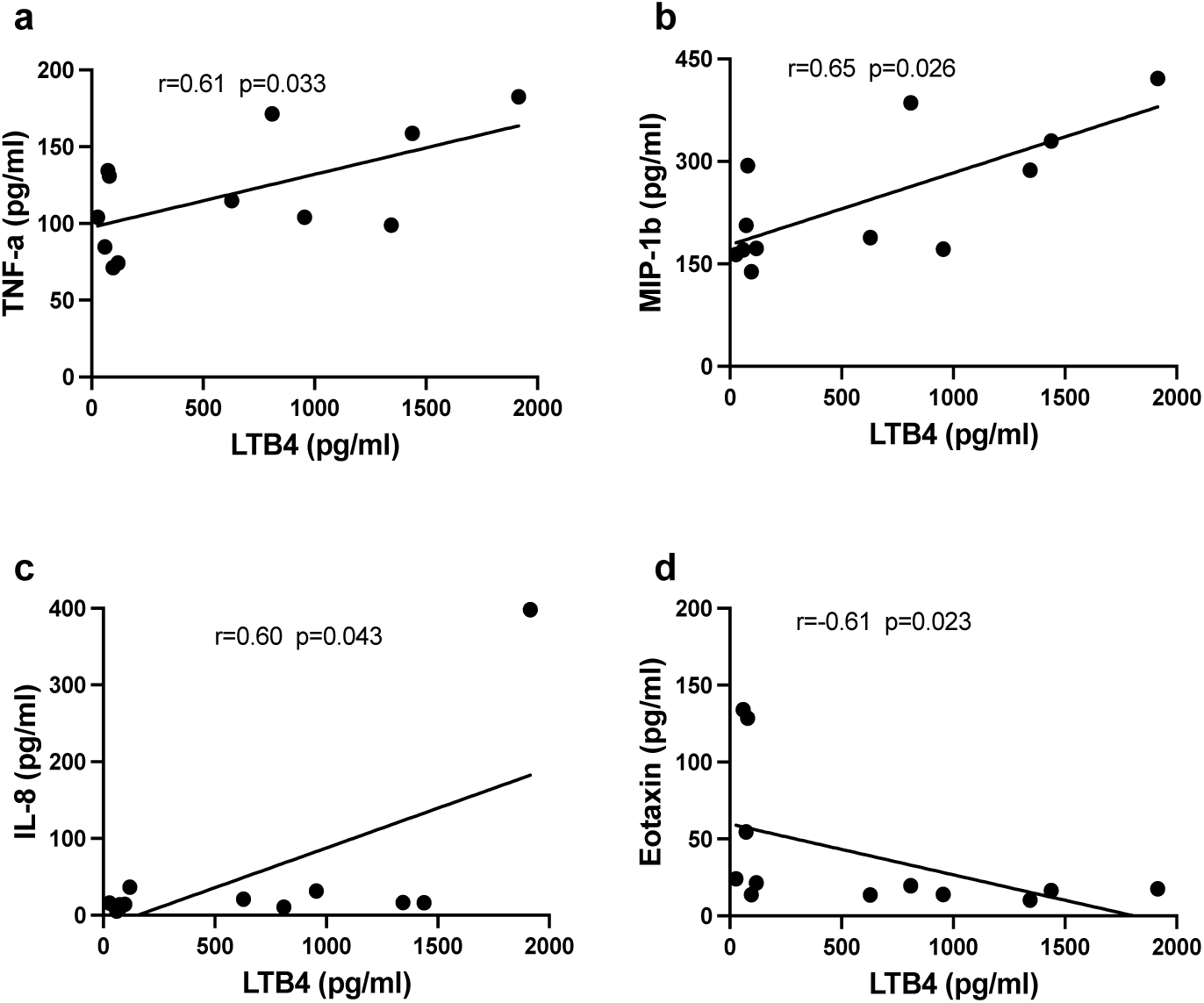
**a - h,** The expression level of LTB4 measured using ELISA, was correlated with circulatory cytokines/chemokines and growth factors including TNF-α, MIP-1b, IL-8, and Eotaxin from matching active TB participants. A Pearson correlation coefficient was used to assess the linear relationship between LTA4H and the inflammatory cytokines.

**Figure S5.**
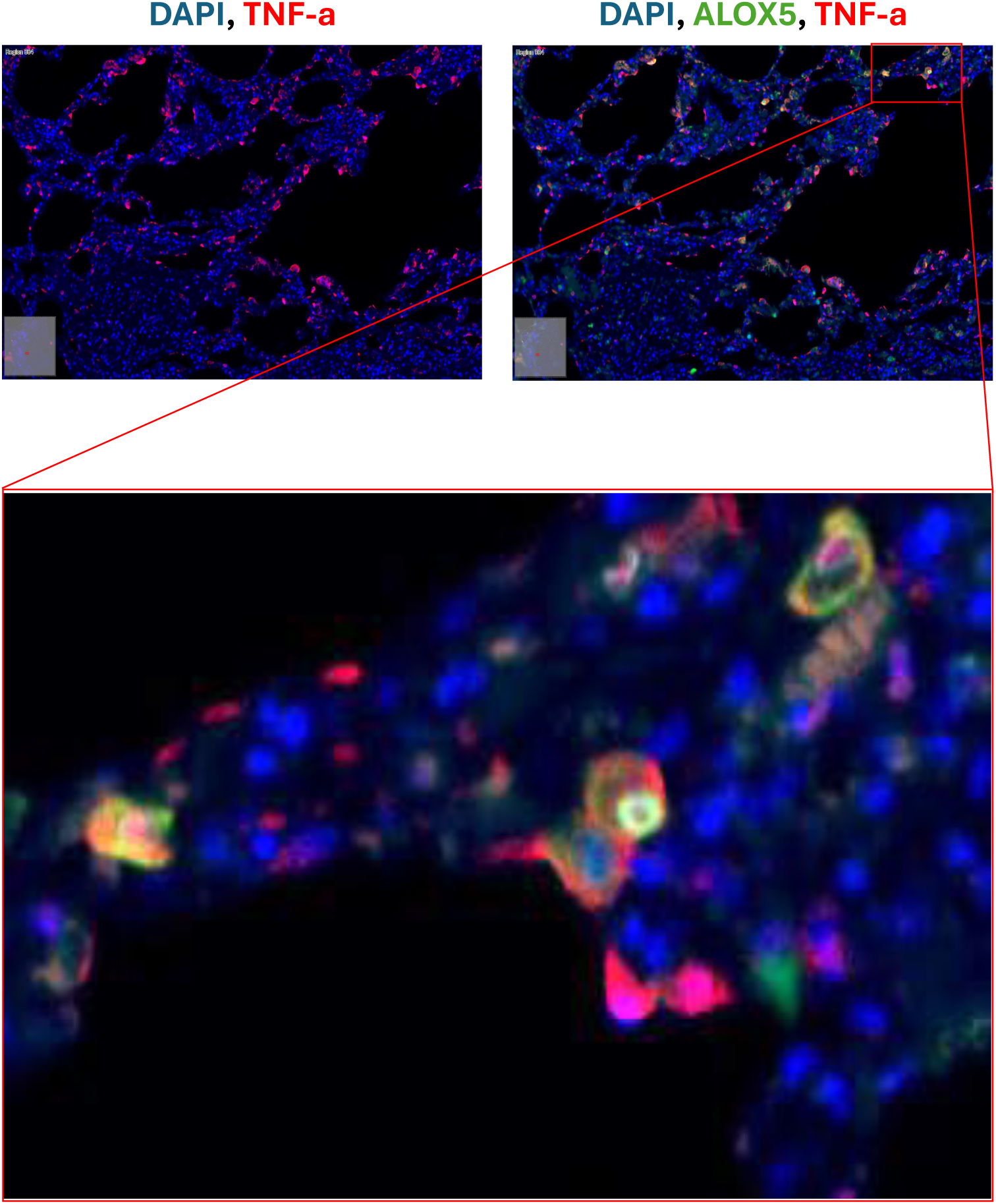
IF staining of ALOX5 and TNF-α to demonstrate their co-localization within the same cell populations obtained from TB diseased lung.

**Figure S6.**
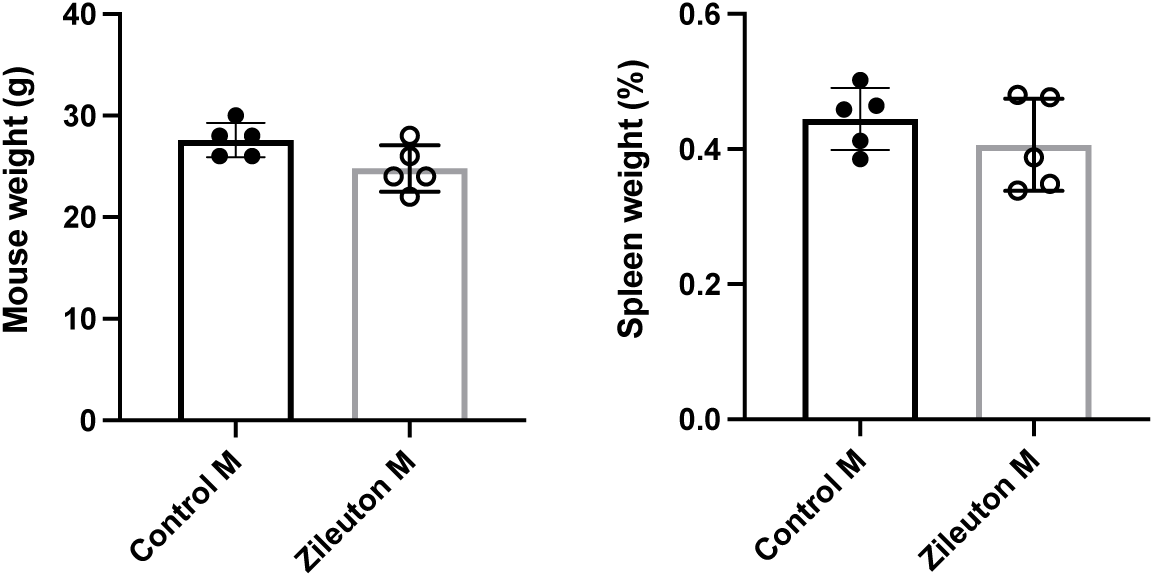
Lung tissue weights were measured from treated and non-treated mice and compared at day 56 post infection.

## Graphical Abstract 1

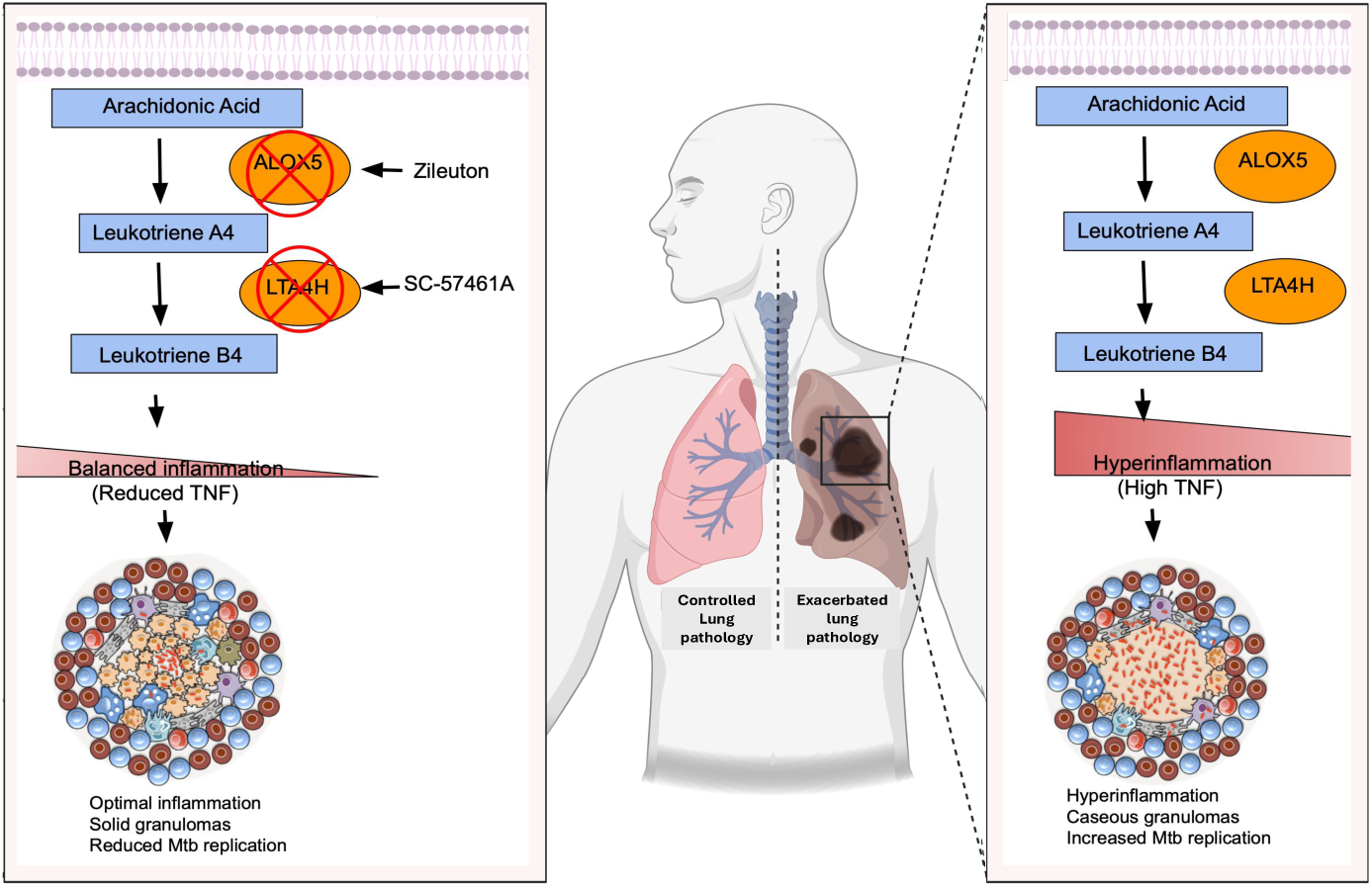

## Notes

### Competing Interest Statement

The authors have declared no competing interest.

