## Supplemental Table 2 for "Targeting ALOX5/LTA4H driven granuloma caseation as a host-directed strategy for control of TB associated lung damage"

| **Supplemental Table 2:** Clinical characteristics of study participants for blood samples | | | |
| --- | --- | --- | --- |
| Characteristics | Healthy  (IGRA-) | LTBI  (IGRA+) | TB (GeneXpert+) |
| n (70) | 20 | 20 | 30 |
| **Age (mean and SD)** | 30.90± 12.59 | 32.00±11.95 | 35.37 ± 10.93 |
| **Sex** |  |  |  |
| Male | 35.% (7) | 45% (9) | 73.33 % (22) |
| Female | 65% (13) | 55% (11) | 23.33% (7) |
| Not disclosed |  |  | 3.33% (1) |
| **Assays** |  |  |  |
| plasma - Bio-Plex | 20 | 20 | 30 |
| qPCR on whole blood RNA | 20 | 20 | 30 |
| Plasma ELISA/Calorimetric assays: ALOX5, LTA4H, LTB4 and AA | 20 | 20 | 30 |
